# HAUSP Stabilizes SOX2 through Deubiquitination to Maintain Self-renewal and Tumorigenic Potential of Glioma Stem Cells

**DOI:** 10.1101/2021.06.09.447550

**Authors:** Zhi Huang, Kui Zhai, Qiulian Wu, Xiaoguang Fang, Qian Huang, Weiwei Tao, Justin D. Lathia, Jennifer S. Yu, Jeremy N. Rich, Shideng Bao

## Abstract

Glioblastoma (GBM) is the most lethal brain tumor containing glioma stem cells (GSCs) that promote malignant growth and therapeutic resistance. The self-renewal and tumorigenic potential of GSCs are maintained by core stem cell transcription factors including SOX2. Defining the posttranslational regulation of SOX2 may offer new insights into GSC biology and potential therapeutic opportunity. Here, we discover that HAUSP stabilizes SOX2 through deubiquitination to maintain GSC self-renewal and tumorigenic potential. HAUSP is preferentially expressed in GSCs in perivascular niches in GBMs. Disrupting HAUSP by shRNA or its inhibitor P22077 promoted SOX2 degradation, induced GSC differentiation, impaired GSC tumorigenic potential, and suppressed GBM tumor growth. Importantly, pharmacological inhibition of HAUSP synergized with radiation to inhibit GBM growth and extended animal survival, indicating that targeting HAUSP may overcome GSC-mediated radioresistance. Our findings reveal an unappreciated crucial role of HAUSP in the GSC maintenance and provide a promising target for developing effective anti-GSC therapeutics to improve GBM treatment.

**Highlights:**

1. HAUSP deubiquitinates and stabilizes SOX2 in glioma stem cells (GSCs).
2. HAUSP is preferentially expressed by GSCs in perivascular niches in GBMs.
3. HAUSP is required for maintaining GSC self-renewal and tumorigenic potential.
4. Targeting HAUSP inhibited malignant growth in GSC-derived GBM xenografts.
5. Inhibition of HAUSP synergized with radiation to suppress GBM tumor growth.

## Introduction

Glioblastoma (GBM) is the most malignant and lethal type of primary brain tumor with extremely poor prognosis. Despite maximal therapy and new advances in cancer treatment, the median survival of GBM patients remain approximately 14-16 months ^1–3^. GBMs are highly resistant to current therapies including radiation, chemotherapy, targeted therapy and immune checkpoint inhibition ^4–6^, leading to rapid and essentially universal tumor recurrence after the treatment. Thus, developing novel effective therapeutics is critical for improving the survival of GBM patients. GBMs display striking cellular heterogeneity and hierarchy in differentiation status. The cancer cells at the apex of the cellular hierarchy within a GBM tumor bulk are glioma stem cells (GSCs) or tumor initiating cells that display potentials of extensive self-renewal, multi-lineage differentiation, and propagation of tumors in vivo ^7–15^. It has been well-recognized that GSCs are critical cancer cells to maintain tumor growth and repopulate tumor after current treatments. We and others have demonstrated that GSCs interact with their niches to promote tumor angiogenesis, cancer invasion, immune evasion, and therapeutic resistance ^16–25^. We also discovered that GSCs give rise to the majority of vascular pericytes in GBM ^26^. Importantly, these GSC-derived pericytes contribute to the formation of the blood-brain tumor barrier (BTB) that blocks drug delivery and negatively impacts chemotherapy ^27^. Furthermore, we found that GSCs secrete periostin (POSTN) to recruit monocyte-derived macrophages (TAMs) and secrete WISP1 to maintain the recruited TAMs as tumor-supportive macrophages (M2) that promote GBM malignant progression ^23, 25^. These studies indicate that targeting GSCs may effectively inhibit GBM malignant growth, disrupt the BTB, remodel the tumor microenvironment (TME) including the immune microenvironment, and overcome therapeutic resistance to improve treatment efficacy and reduce tumor recurrence. We aim to develop novel therapeutics through targeting key signaling regulators in GSC maintenance to directly eliminate GSC populations or induce GSC differentiation, which may effectively disrupt GSC tumorigenic potential to improve GBM treatment. Thus, dissecting signaling pathways controlling GSC self-renewal and tumorigenic potential is of highest importance.

As GSCs share critical characteristics with neural stem cells and embryonic stem cells (ESCs), some crucial stem cell transcription factors (SCTFs) including SOX2 involved in the maintenance of normal stem cells are also required for maintaining GSC properties. As one of key components of the SCTF network, SOX2 plays critical roles in maintaining the self-renewal potential of normal stem cells and cancer stem cells (CSCs), including GSCs ^28–30^. Disrupting SOX2 with shRNA in GSCs potently induces cell differentiation. SOX2 is an important member of the SRY-related HMG-box protein family and contains the DNA-binding domain, the transcriptional activation domain, and the group B homolog domain ^31^. It has been shown that SOX2 is highly expressed in several types of malignant tumors ^32–35^. Importantly, SOX2 is preferentially expressed in CSC populations, including GSCs in GBM, relative to their matched differentiated progeny ^29, 36^. As a core stem cell transcription factor, SOX2 is tightly and dynamically regulated at both transcriptional and posttranslational levels to affect cellular states during normal development. In mouse ESCs, the SOX2 protein is regulated by the ubiquitination mediated by WWP2, an ubiquitin E3 ligase that targets SOX2 for degradation ^37^. However, how SOX2 is regulated by ubiquitination and deubiquitination in GSCs remains elusive. As SOX2 plays a vital role in maintaining GSCs that are critical cells for promoting malignant progression and therapeutic resistance, it is important to define the posttranslational regulation of SOX2 in GSCs, which may provide useful molecular targets for developing novel therapeutics targeting GSCs to overcome GSC-driven therapeutic resistance and improve GBM treatment.

In this study, we aim to identify the key potential deubiquitinase that stabilizes SOX2 to maintain GSC self-renewal and tumorigenic potential. To this end, we found that HAUSP (the Herpesvirus-Associated Ubiquitin-Specific Protease, also known as Ubiquitin-Specific Protease 7, USP7) is the critical deubiquitinase that stabilizes SOX2 protein through deubiquitination to promote the GSC maintenance. HAUSP was initially identified as an ubiquitin-specific protease (USP) associated with the viral protein ICP0 (herpes simplex virus type 1 regulatory protein) or EBNA1 (Epstein-Barr nuclear antigen 1) during viral infection, thus regulating ICP0 stability as well as EBNA1 transcriptional activity ^38, 39^. HAUSP plays vital roles in regulating critical signaling pathways in oncogenesis ^40–46^. This deubiquitinase has been shown to be overexpressed in breast carcinoma ^47^, lung squamous cell carcinoma and large cell carcinoma ^48^, chronic lymphocytic leukemia ^49^, prostate cancer ^44^, and epithelial ovarian cancer ^50, 51^. HAUSP is well-known for its ability to stabilize the oncogenic protein MDM2 (or the human ortholog HDM2) that functions as an ubiquitin E3 ligase to target p53 for proteasome-mediated degradation through ubiquitination ^40, 42, 52^. Thus, HAUSP is a negative regulator of the tumor suppressor p53 under normal conditions. Genetic ablation of HAUSP results in reduced MDM2 protein levels, leading to stabilization and accumulation of p53 and growth suppression of cancer cells ^40, 42, 52^. HAUSP also regulates the stability and function of several other critical proteins such as PTEN, FOX4, Claspin, DNMT1 and REST under normal and stress conditions ^41, 44, 53–55^. For example, HAUSP is able to deubiquitinate the tumor suppressor PTEN, and thus regulates its nuclear exclusion ^44^. In response to oxidative stress, HAUSP inhibits transcriptional activity and nuclear localization of FOXO4 (a forkhead box O transcription factor) by interacting with and deubiquitinating FOXO protein ^54^. During early embryonic development, HAUSP deletion caused early embryonic lethality between E6.5 and E7.5, which was thought due to increased p53 levels to arrest cell cycle ^56, 57^. Interestingly, HAUSP and p53 double knockout mice, although extending the embryonic development, could not survive the whole prenatal development ^56, 57^. Inactivation of p53 failed to completely rescue the neonatal lethality of hausp-null mice, suggesting a p53-independent function of HAUSP in the control of cell proliferation or in the maintenance of stem cells. However, the potential roles of HAUSP in the maintenance of stem cells including cancer stem cells such as GSCs have not been explored.

Our study showed that HAUSP is preferentially expressed in GSCs relative to matched non-stem tumor cells (NSTCs) in GBMs. We demonstrated that HAUSP functions as a critical deubiquitinase that stabilizes SOX2 protein through deubiquitination to maintain GSC phenotype and tumorigenic potential. Disruption or functional inhibition of HAUSP by shRNA or its inhibitor P22077 promoted SOX2 degradation, induced GSC differentiation, impaired tumorigenic potential of GSCs, and suppressed GBM malignant growth. Importantly, pharmacological inhibition of HAUSP synergized with radiation to potently inhibit GBM tumor growth and extended animal survival, indicating that disrupting GSCs by targeting HAUSP may overcome GBM radioresistance. Our findings not only reveal an unappreciated crucial role of HAUSP in maintaining GSC self-renewal and tumorigenic potential but offer an attractive functional target for developing effective therapeutics targeting GSC populations to overcome therapeutic resistance, which may effectively improve the survival of GBM patients.

## Results

### HAUSP deubiquitinates and stabilizes SOX2 in glioma stem cells (GSCs)

To identify the potential deubiquitinase(s) responsible for SOX2 deubiquitination and stabilization in GSCs, we analyzed SOX2-interacting proteins through mass spectrometric analyses to determine deubiquitinase(s) associated with SOX2 in GSCs. We found that one important deubiquitinase called HAUSP (USP7) was present only in the immunoprecipitation (IP) complex pulled down by anti-Flag-SOX2 in GSCs (Figure S1A). Further IP and immunoblot analyses confirmed the interaction between HAUSP and the endogenous SOX2 or ectopic Flag-SOX2 in GSCs (Figures 1A and 1B). The ubiquitination assay demonstrated that HAUSP disruption by shRNA (shHAUSP) or functional inhibition by its inhibitor P22077 ^58, 59^ markedly increased SOX2 ubiquitination in GSCs (Figures 1C and S1B), indicating that HAUSP is the key deubiquitinase of SOX2 in GSCs. Immunoblot and immunofluorescent staining validated that disrupting HAUSP by shRNA in GSCs dramatically reduced SOX2 protein levels (Figures 1D and 1E), but a short-term disruption of HASUP did not significantly affect mRNA levels of SOX2 in GSCs (Figure S1C), indicating that HAUSP stabilizes SOX2 protein in GSCs. Moreover, ectopic expression of the wild-type HAUSP (HAUSP-Wt) but not the catalytically dead HAUSP mutant (HAUSP-CS) reduced ubiquitination of SOX2 and increased SOX2 protein levels (Figure 1F). Furthermore, in vitro ubiquitination assay validated that wild-type HASUP (Flag-HAUSP-Wt) but not the catalytically dead HAUSP (Flag-HAUSP-CS) was able to deubiquitinate and stabilize SOX2 proteins (HA-SOX2) (Figure S1D). Collectively, these data demonstrate that HAUSP is the key deubiquitinase that stabilizes SOX2 through deubiquitination in GSCs, indicating that HAUSP plays an essential role in maintaining SOX2 protein levels in GSCs.

**Figure 1.**
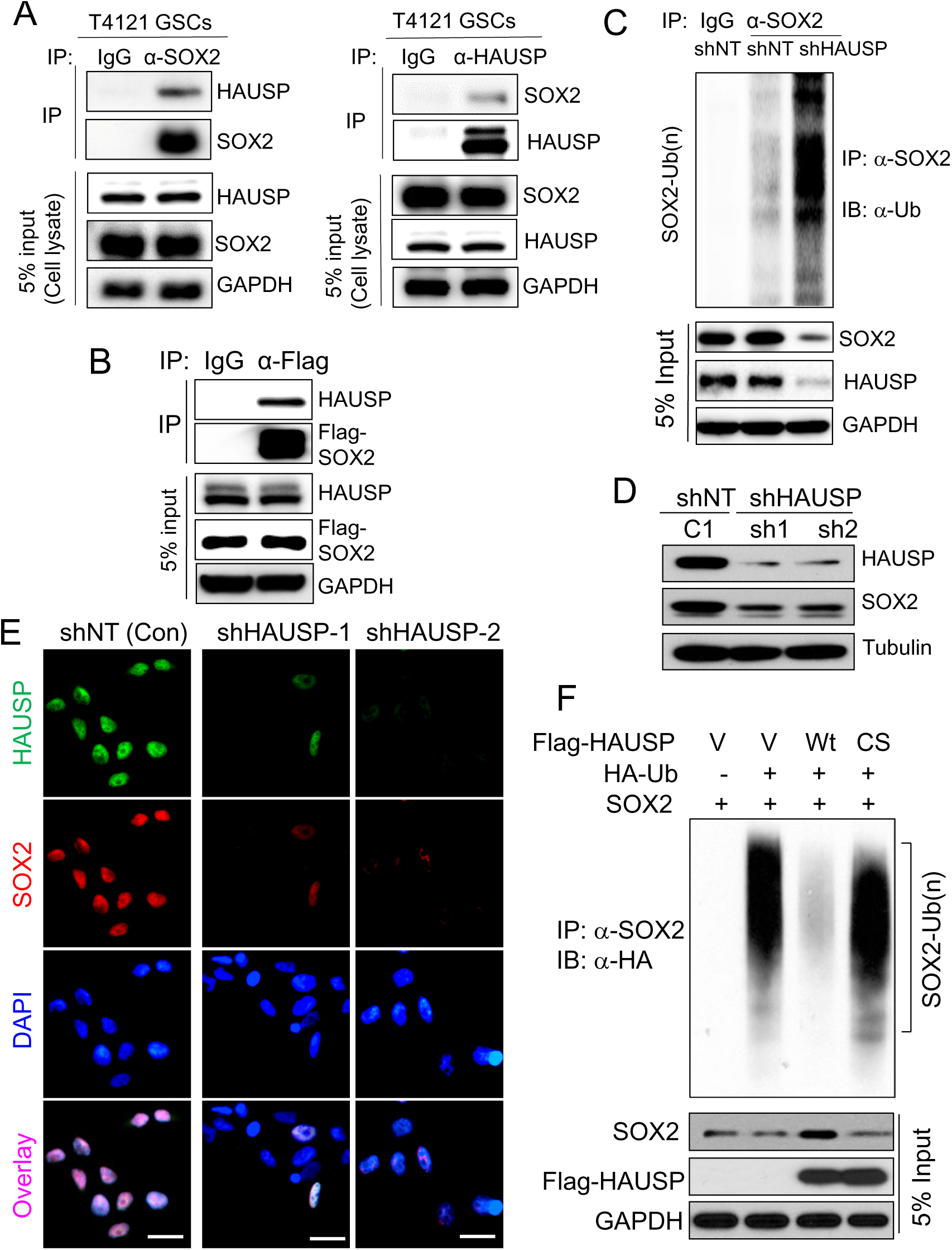
HAUSP interacts with SOX2 and mediates deubiquitination of SOX2 to stabilize SOX2 in glioma stem cells (GSCs). **(A)** Co-immunoprecipitation (Co-IP) to validate the interaction between endogenous SOX2 and HAUSP in GSCs. Cell lysates of GSCs (T4121) were co-immunoprecipitated with anti-SOX2 or anti-HAUSP specific antibodies. The Co-IP products and total cell lysates were immunoblotted with anti-SOX2 or anti-HAUSP antibody. **(B)** Co-immunoprecipitation (Co-IP) to confirm the interaction between ectopically expressed Flag-SOX2 and endogenous HAUSP in GSCs. GSCs (CCF3264) were transduced with Flag-SOX2 through lentiviral infection, and the cell lysates were co-immunoprecipitated with anti-Flag antibody. The Co-IP products and 5% cell lysates were immunoblotted with anti-Flag and anti-HAUSP antibodies. **(C)** Ubiquitination assay to examine the effect of HAUSP knockdown on poly-ubiquitination of SOX2 in GSCs. T4121 GSCs were transduced with HAUSP shRNAs (shHAUSP) or non-targeting shRNA (shNT) through lentiviral infection for 48 hours, then treated with MG132 for 8 hours, and harvested for immunoprecipitation (IP) with anti-SOX2 antibody or IgG control. The IP products were immunoblotted with anti-ubiquitin (Ub) antibody to assess the SOX2 ubiquitination in GSCs with or without HAUSP disruption. 5% cell lysates were immunoblotted with antibodies against SOX2, HAUSP and GAPDH (control). **(D)** Immunoblot analyses of SOX2 and HAUSP in GSCs transduced with shHAUSP or shNT. GSCs (T4121) were transduced with two independent HAUSP shRNAs (shHAUSP, sh1 or sh2) or non-targeting shRNA (shNT, C1) through lentiviral infection for 48 hours, and then harvested for immunoblot analyses with specific antibodies against SOX2, HAUSP and GAPDH (control). **(E)** Immunofluorescent staining of SOX2 and HAUSP to validate the effect of HAUSP knockdown on SOX2 protein levels in GSCs. T4121 GSCs were transduced with two independent HAUSP shRNAs (shHAUSP-1 or shHAUSP-2) or non-targeting shRNA (shNT) through lentiviral infection for 48 hours, then immunostained with specific antibodies against SOX2 (in green) and HAUSP (in red), and counterstained with DAPI to indicate nuclei (in blue). Scale bar, 20 µm. **(F)** Ubiquitination assay to examine the effect of overexpressed wild-type HAUSP (Wt) or catalytically dead mutant HAUSP (CS) on poly-ubiquitination of SOX2 in GSCs. T4121 GSCs were transduced with wild-type Flag-HAUSP (Wt), catalytically dead Flag-HAUSP (CS), or vector control through lentiviral infection for 48 hours, then treated with MG132 for 8 hours, and harvested for immunoprecipitation (IP) with anti-SOX2 antibody or IgG control. The IP products were immunoblotted with anti-ubiquitin (Ub) antibody to assess the SOX2 ubiquitination in GSCs overexpressing wild-type HAUSP or the mutant HAUSP. 5% cell lysates were immunoblotted with antibodies against SOX2, Flag-HAUSP and GAPDH (control).

### HAUSP is preferentially expressed by GSC populations

As SOX2 is preferentially expressed by GSCs relative to matched non-stem tumor cells (NSTCs), and SOX2 functions as an essential stem cell transcription factor for the GSC maintenance, we next examined whether its deubiquitinase HAUSP is also differentially expressed by GSCs relative to NSTCs. GSCs and matched NSTCs were isolated from primary GBM tumors or patient-derived xenografts (PDXs) and validated by functional assays including limiting dilution tumorsphere formation, in vitro differentiation, and in vivo tumor formation ^16, 23, 26, 27, 60–62^. Immunoblot analysis demonstrated that much higher levels of HAUSP were expressed by GSC populations expressing SOX2 and OLIG2 relative to matched NSTCs isolated from several GBM surgical specimens or PDX tumors (Figures 2A and S2A). Quantitative RT-PCR analyses showed that GSCs expressed much higher levels of HAUSP mRNA levels than the matched NSTCs (Figures 2B and S2B), indicating that up-regulation of HAUSP in GSCs occurs at the transcriptional level. Immunofluorescent staining further confirmed that HAUSP and the GSC marker SOX2 or OLIG2 are preferentially expressed by GSCs and co-localized in nuclei of GSCs relative to matched NSTCs (Figures 1E, 2C and S2C). Consistently, HAUSP is preferentially expressed by GSCs co-expressing SOX2 in total glioma cells freshly isolated from primary GBMs (Figure 2D). Moreover, GSC-derived tumorspheres maintain high levels of HAUSP expression in cells co-expressing SOX2 or OLIG2 as demonstrated by immunofluorescent staining (Figures 2E and S2D). Collectively, these data demonstrate that HAUSP is preferentially expressed by GSC populations isolated from primary GBMs and PDXs.

**Figure 2.**
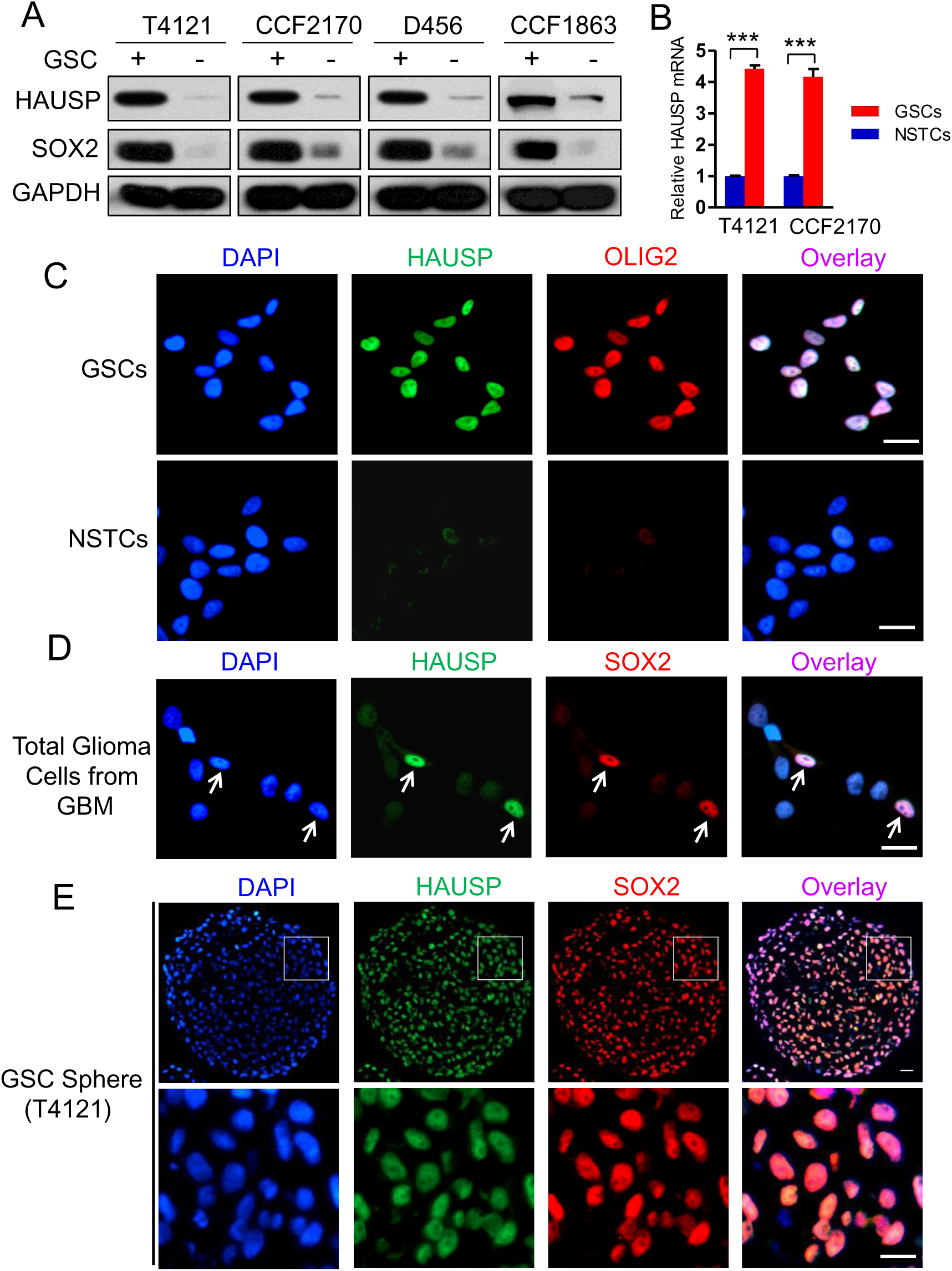
HAUSP is preferentially expressed by glioma stem cells (GSCs) isolated from human primary GBMs. **(A)** Immunoblot analyses of HAUSP and SOX2 in GSCs and matched non-stem tumor cells (NSTCs) isolated from four GBM tumors. HAUSP is preferentially expressed in GSCs expressing high levels of SOX2 relative to NSTCs expressing low levels of SOX2. **(B)** Quantitative RT-PCR analyses of HAUSP mRNA expression in GSCs and matched NSTCs. HAUSP mRNA is differentially expressed in GSCs relative to NSTCs. Student t test was used to assess the significance. ***, p<0.001. **(C)** Immunofluorescent staining of HAUSP and OLIG2 in GSCs and matched NSTCs isolated from primary GBMs. GSCs and matched NSTCs were immunostained with specific antibodies against HAUSP (in green) and SOX2 (in red), and then counterstained with DAPI to indicate nuclei (in blue). Scale bar, 20 µm. **(D)** Immunofluorescent staining of HAUSP and SOX2 in total glioma cells isolated from a primary GBM. Total glioma cells from a GBM (CCF2045) were immunostained with specific antibodies against HAUSP (in green) and SOX2 (in red), and then counterstained with DAPI to indicate nuclei (in blue). HAUSP is preferentially expressed in the cells expressing the GSC marker SOX2 (indicated by white arrows). Scale bar, 20 µm. **(E)** Immunofluorescent staining of HAUSP and SOX2 in GSC tumorsphere. Sections of GSC tumorspheres (T4121) were immunostained with specific antibodies against HAUSP (in green) and SOX2 (in red), and then counterstained with DAPI to indicate nuclei (in blue). A small portion of the GSC sphere indicated by the white square was enlarged and shown in the bottom panels. Scale bar, 15 µm.

### HAUSP is highly expressed by GSCs in perivascular niches in human GBMs

To validate the preferential expression of HAUSP by GSCs in human GBM tumors in vivo, we examined the expression patterns of HAUSP and the GSC marker SOX2 or OLIG2 in several GBM surgical specimens. Double immunofluorescent staining showed that HAUSP is specifically expressed by a subpopulation of glioma cells expressing SOX2 or OLIG2, two well-established GSC markers, in human primary GBM tumors (Figure 3A), supporting that HAUSP is differentially expressed by GSCs relative to NSTCs in human GBMs. Interestingly, those GSCs expressing HAUSP are often localized in the area near blood vessels marked by CD31 (Figures 3B and S3A), which is consistent with our previous finding that GSCs are preferentially distributed in perivascular niches in GBMs ^17, 23, 26, 62^. In addition, immunohistochemical (IHC) staining further confirmed that HAUSP is mainly localized in nuclei in a fraction of glioma cells near blood vessels in primary GBM tumors (Figures 3C and S3B). In contrast, HAUSP is not detectable in the normal brain tissues (Figures 3C and S3C). These data demonstrate that HAUSP is preferentially expressed by GSCs in perivascular niches in human GBM tumors, indicating that HAUSP is a potential marker to identify GSC populations in human GBM tumors.

**Figure 3.**
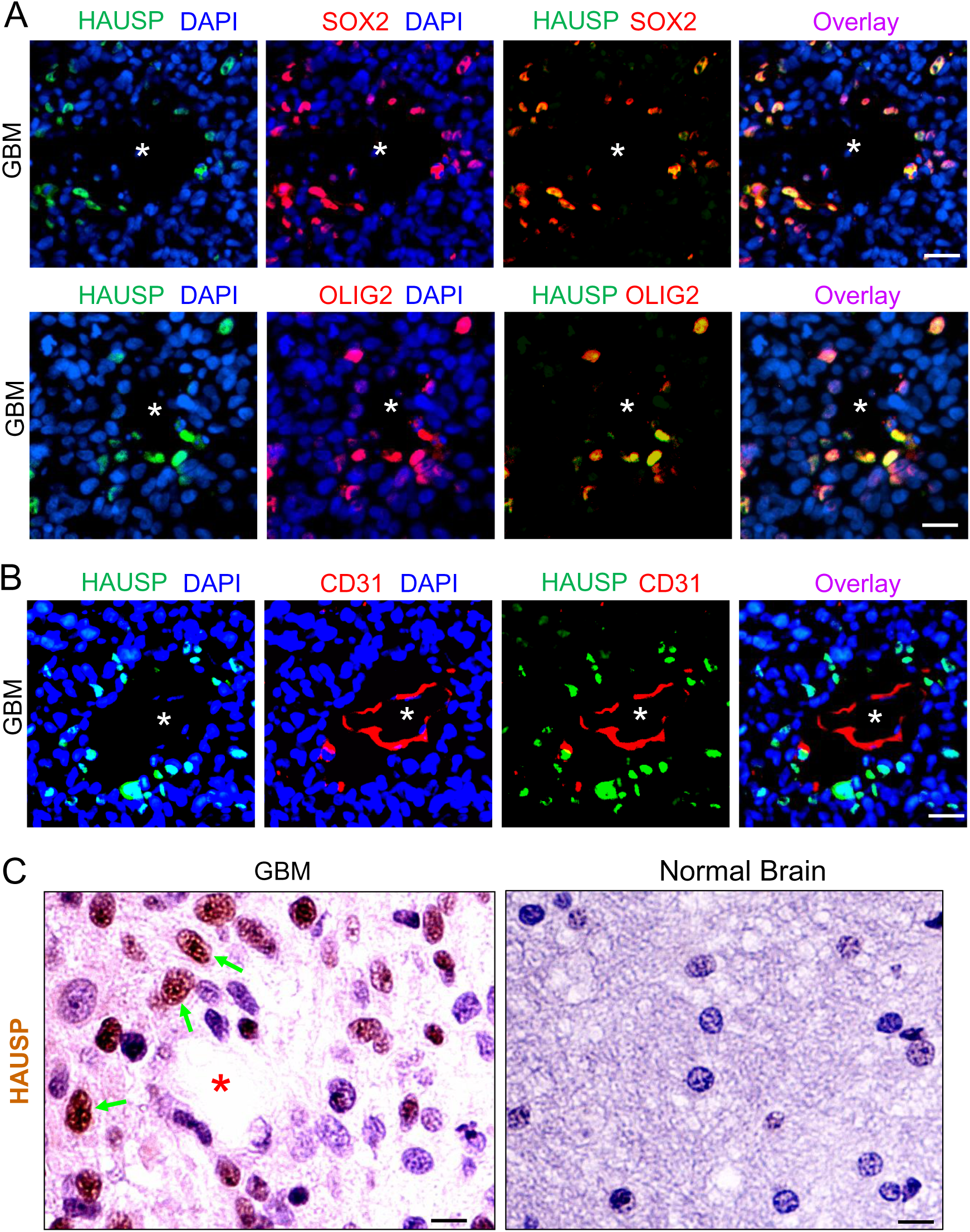
HAUSP and SOX2 are co-expressed in GSCs in perivascular niches in human primary GBMs. **(A)** Immunofluorescent staining of HAUSP and SOX2 or OLIG2 in human primary GBM. Frozen sections of GBM (CCF1992) were immunostained with specific antibodies against HAUSP (in green) and SOX2 or OLIG2 (in red), and then counterstained with DAPI to indicate nuclei (in blue). White stars indicate potential vessel lumens in the GBM tumor. Scale bar, 25 µm. **(B)** Immunofluorescent staining of HAUSP and the endothelial cell marker CD31 in human primary GBMs. Frozen sections of GBM were immunostained with specific antibodies against HAUSP (in green) and CD31 (in red), and then counterstained with DAPI to indicate nuclei (in blue). White stars indicate the vessel lumen in the GBM tumor. Scale bar, 25 µm. **(C)** Immunohistochemical (IHC) staining of HAUSP in human primary GBMs. Paraffin sections of a primary GBM and normal brain tissues were immunostained with a specific anti-HAUSP antibody, and counterstained with hematoxylin to indicate nuclei. HAUSP (in brown) is expressed by some glioma cells around the vessel (indicated by a red star), while HAUSP is not detectable in the normal brain tissue. Scale bar, 10 µm.

### Disruption or inhibition of HAUSP promotes GSC differentiation

As disrupting HAUSP by shRNA reduced SOX2 protein, one of key stem cell transcription factors critically required for the GSC maintenance, we next examined whether disruption or inhibition of HAUSP could impair GSC maintenance to promote GSC differentiation. Immunoblot analysis showed that HAUSP knockdown in GSCs promoted expression of differentiated cell markers such as TUJ1 (a marker for neuronal lineage) and GFAP (a marker for astrocytes) in the in vitro differentiation assay (Figure 4A). RT-PCR analyses confirmed that HAUSP knockdown induced expression of the differentiated cell markers including TUJ1 and GFAP (Figure 4B). Immunofluorescent staining validated that disrupting HAUSP by shRNA markedly increased TUJ1^+^ or GFAP^+^ cells populations (Figures 4C-4F). These data demonstrate that disrupting HAUSP effectively promoted GSC differentiation. Consistently, during GSC differentiation induced by addition of serum or depletion of EGF/bFGF in the culture medium, protein levels of HAUSP and the GSC marker including SOX2 gradually decreased as demonstrated by immunoblotting and immunofluorescent staining (Figures S4A and S4B). Ubiquitination assay demonstrated that SOX2 ubiquitination gradually increased during GSC differentiation (Figure S4C). Furthermore, inhibiting HAUSP by its inhibitor P22077 also promoted expression of the differentiated cell markers including MAP2 (a neuronal marker) and GFAP (an astrocyte marker) but reduced SOX2 levels in a dose- and time-dependent manner (Figures S4D and S4E). Immunofluorescent staining confirmed that P22077 treatment reduced SOX2 protein (Figure S4F) but induced expression of differentiation markers including TUJ1 and GFAP (Figure S4G). Collectively, these data demonstrate that HAUSP plays a critical role in preventing GSC differentiation, indicating that HAUSP is required for the maintenance of GSC stemness. Thus, disrupting or inhibiting HAUSP is an attractive strategy to induce GSC differentiation, which may impair GSC-driven tumor growth and overcome GSC-mediated therapeutic resistance.

**Figure 4.**
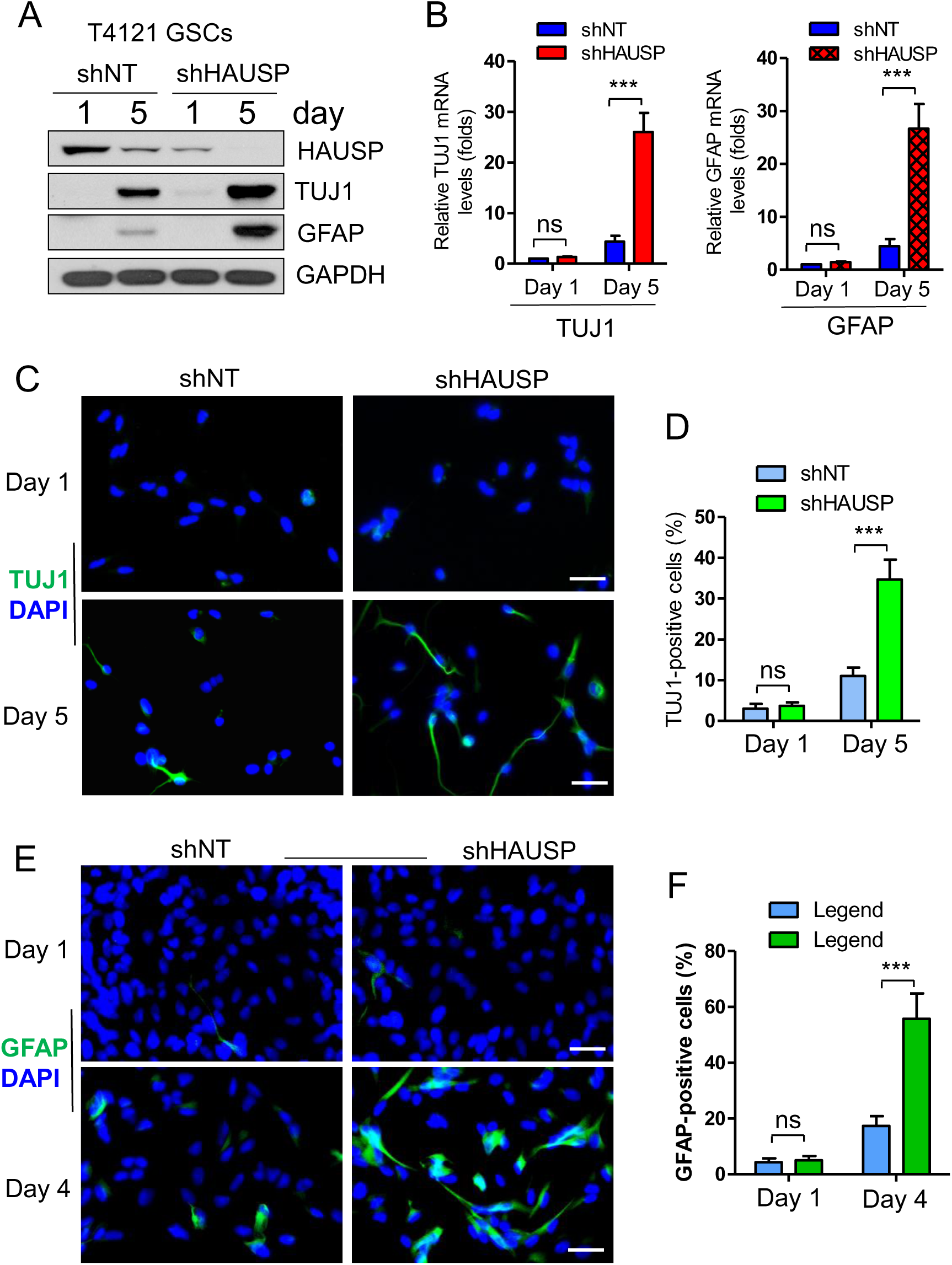
Disrupting HAUSP promoted GSC differentiation in vitro. **(A)** Immunoblot analyses of differentiated cell markers TUJ1 (a neuronal marker) and GFAP (an astrocyte marker) in GSCs expressing shHAUSP or shNT (control). GSCs (T4121) were transduced with shHAUSP or shNT through lentiviral infection for 48 hours, then cultured in the medium with reduced growth factors for indicated days, and harvested for immunoblot analyses with specific antibodies against TUJ1, GFAP, HAUSP, and GAPDH (loading control). **(B)** Quantitative PCR (RT-PCR) analyses of TUJ1 and GFAP mRNA levels in GSCs expressing shHAUSP or shNT (control). GSCs (T4121) were transduced with shHAUSP or shNT through lentiviral infection for 48 hours, then cultured in the medium with reduced growth factors for the indicated days, and harvested for RT-PCR analyses. Student t test was used to assess the significance. ***, p<0.001. (**C** and **D**) Immunofluorescent staining of TUJ1 (a neuronal marker) in GSCs expressing shHAUSP or shNT (control). GSCs (T4121) were transduced with shHAUSP or shNT through lentiviral infection for 48 hours, then cultured in the medium with reduced growth factors for indicated days, and immunostained with a specific antibody against TUJ1 (in green) and counterstained with DAPI to indicate nuclei (in blue). Scale bar, 30 µm. Quantifications of TUJ1-positive cells were shown (D). Student t test was used to assess the significance. ns, no significant difference; ***, p<0.001. (**E** and **F**) Immunofluorescent staining of GFAP (an astrocyte marker) in GSCs expressing shHAUSP or shNT (control). GSCs (T4121) were transduced with shHAUSP or shNT through lentiviral infection for 48 hours, then cultured in the medium with reduced growth factors for indicated days, and immunostained with a specific antibody against GFAP (in green) and counterstained with DAPI to indicate nuclei (in blue). Scale bar, 30 µm. Quantifications of GFAP-positive cells were shown (F). Student t test was used to assess the significance. ns, no significant difference; ***, p<0.001.

### Disrupting HAUSP impairs GSC proliferation and inhibits GBM tumor growth

Because disruption or inhibition of HAUSP promoted GSC differentiation, we next examined whether HAUSP is required for maintaining the proliferation potential of GSCs. Disrupting HAUSP expression by two independent shRNAs (shHAUSP-1 and shHAUSP-2) in GSCs markedly reduced HAUSP protein levels (Figure S5A), resulting in a dramatic inhibition of GSC tumorsphere formation as both sphere number and size were significantly reduced by HAUSP disruption (Figures S5B-S5D). As HAUSP is preferentially expressed in GSCs relative to matched NSTCs, we then examined whether disrupting HAUSP impacts cell growth of GSCs and NSTCs. Cell viability assay showed that silencing HAUSP by two specific shRNAs potently inhibited cell growth of GSCs but showed little effect on growth of matched NSTCs (Figures S5E and S5F). Collectively, these data indicate that HAUSP is critical for maintaining the proliferation potential of GSCs. Thus, targeting HAUSP may represent a potential therapeutic approach to suppress GSC-driven tumor growth.

As GSCs are critical cancer cells to maintain GBM tumor growth, and disrupting HAUSP not only promoted GSC differentiation but also impaired GSC proliferation potential, we next examined the effect of disrupting HAUSP on GSC-driven tumor growth. GSCs expressing luciferase and HAUSP shRNA (shHAUSP) or non-targeting shRNA (shNT) were transplanted into immunocompromised mice through intracranial injection. Bioluminescent imaging of intracranial tumor growth demonstrated that disrupting HAUSP by shRNA potently suppressed GBM tumor growth in GSC-derived xenografts (Figures 5A and 5B). Necropsy of animals sacrificed simultaneously after GSC transplantation confirmed that disrupting HAUSP markedly reduced tumor growth in mouse brains (Figure 5C). As a consequence, the group of mice bearing GBM tumors derived from the shHAUSP-expressing GSCs displayed a significant longer survival than the control group of mice bearing the tumors derived from GSCs expressing shNT (Figures 5D and 5E). Further immunofluorescent analyses indicated that GSC populations (SOX2+ or OLIG2+) were significantly reduced but the differentiated cells (GFAP+) were markedly elevated in those tumors expressing shHAUSP relative to the control tumors expressing shNT (Figures 6A-6D). Moreover, those tumors expressing shHAUSP had a remarkable reduction in cell proliferation as indicated by Ki67 staining (Figures 6E and 6F) and a significant increase in cell apoptosis as demonstrated by immunofluorescent staining of cleaved caspase 3 (Figures 6G and 6H). Taken together, these data demonstrate that disrupting HAUSP potently induces GSCs differentiation and suppresses GBM tumor growth *in vivo* and thus significantly increases the survival of mice bearing the GBM xenografts.

**Figure 5.**
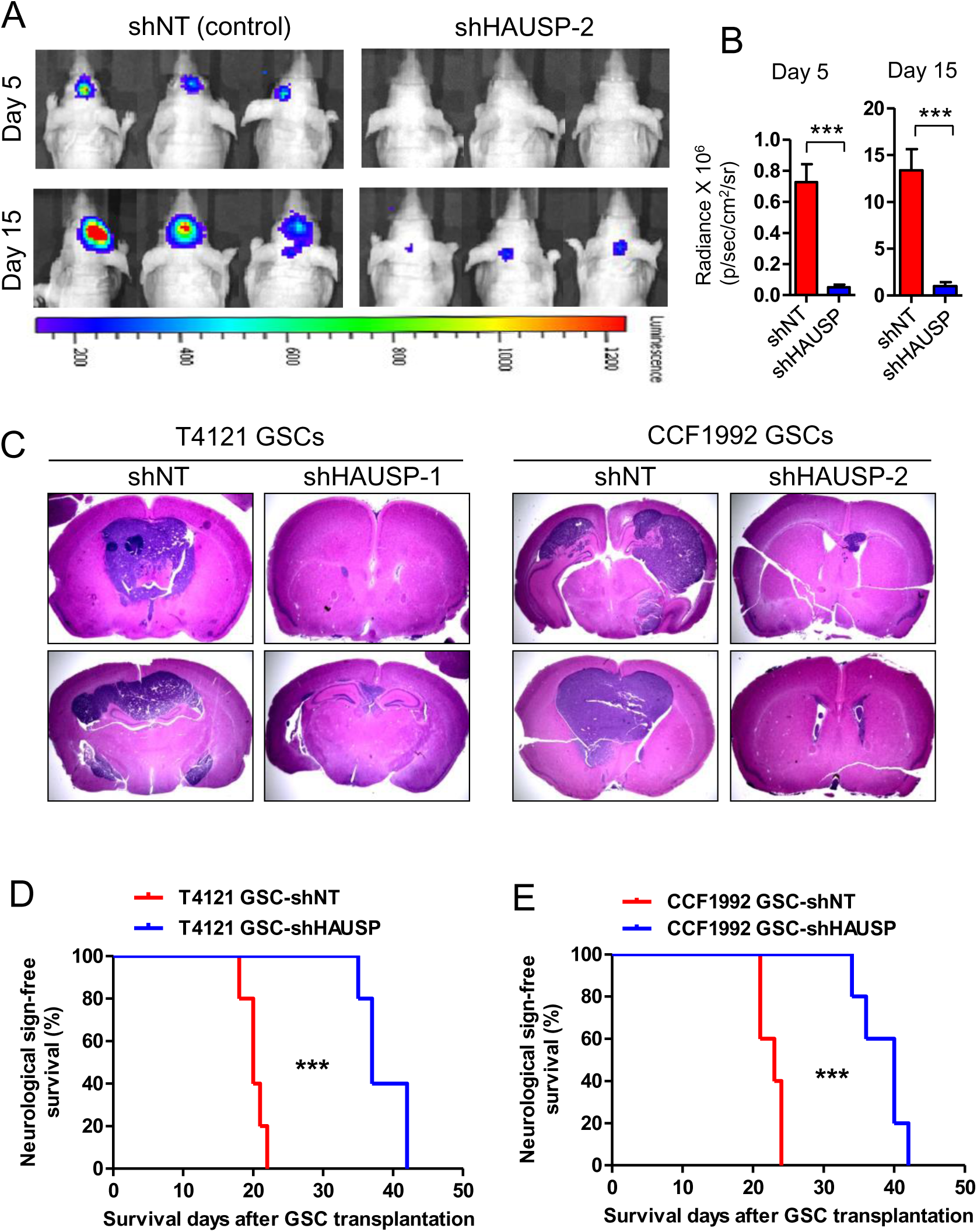
Disrupting HAUSP inhibited GBM tumor growth in GSC-derived xenografts. (**A** and **B**) In vivo bioluminescent imaging (IVIS) of orthotopic GBM xenografts derived from GSCs expressing shHAUSP or shNT (control). GSCs (T4121 or CCF1992) were transduced with luciferase and shHAUSP or shNT through lentiviral infection and then transplanted into brains of immunocompromised nude mice. Intracranial tumor growth was monitored by IVIS. Representative IVIS images at indicated days were shown (A). Quantifications of luciferase intensity on day 5 and day 15 after GSC transplantation were shown (B). Student t test was used to assess the significance. ***, p<0.001. **(C)** Representative images of H&E (hematoxylin and eosin) staining of cross sections of mouse brains bearing GBM xenografts derived from GSCs (T4121 or CCF1992) expressing shRNA targeting HAUSP (shHAUSP) or non-targeting shRNA (shNT). Mouse brains bearing the GBM tumors were harvested on day 21 after the GSC transplantation. (**D** and **E**) Kaplan-Meier survival curves of mice intracranially transplanted with GSCs (T4121 or CCF1992) expressing shRNA targeting HAUSP (shHAUSP) or non-targeting shRNA (shNT). *n*=5 mice. Log-rank analysis was used. **, p<0.01.

**Figure 6.**
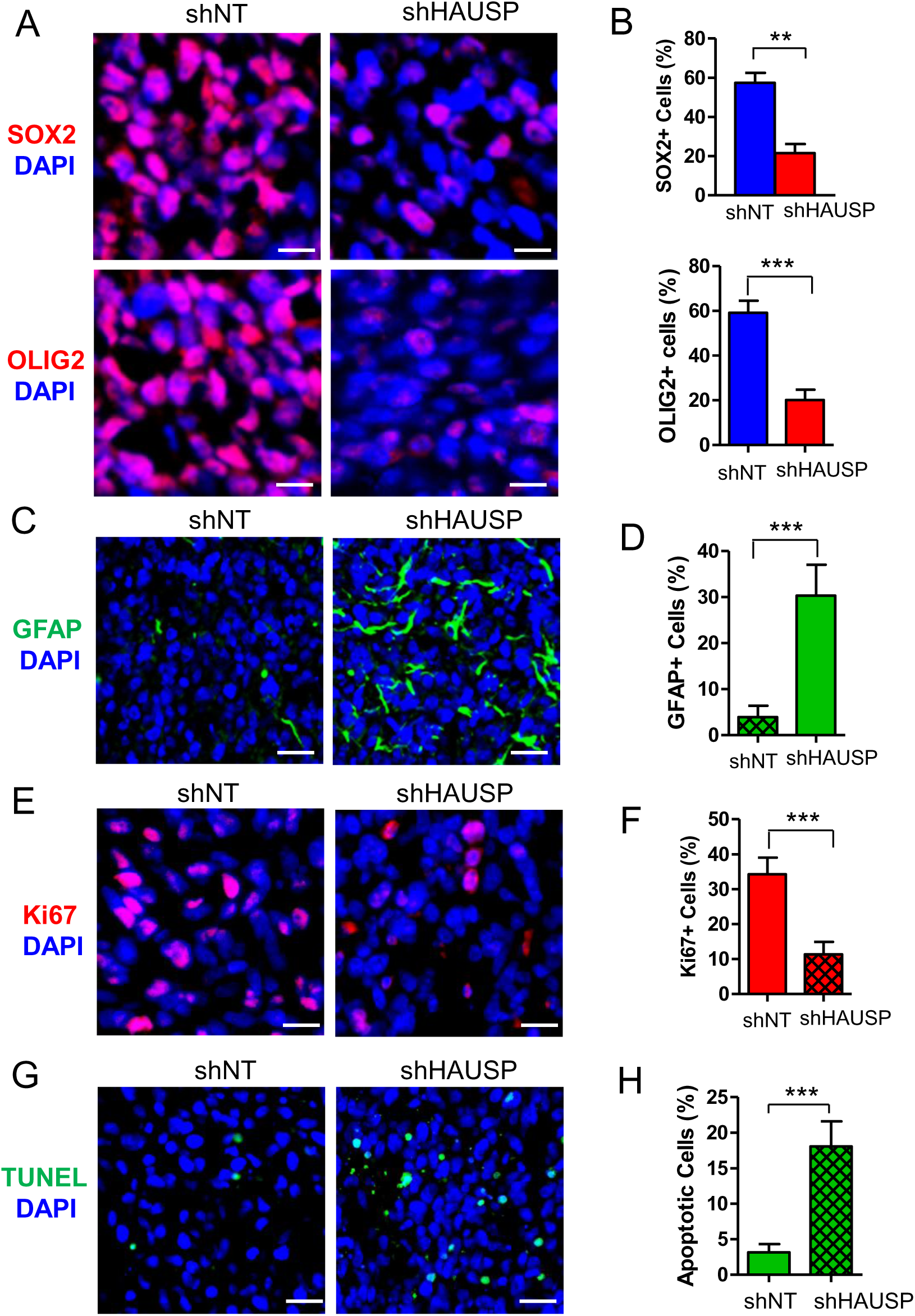
Disrupting HAUSP by shRNA reduced GSC population and cell proliferation, and promoted cell differentiation and apoptosis in vivo. (**A** and **B**) Immunofluorescent staining of the GSC markers SOX2 and OLIG2 in GBM xenografts derived from GSCs (T387) expressing shHAUSP or shNT (control). Tumor sections were immunostained with a specific antibody against SOX2 (in red) or OLIG2 (in red) and counterstained with DAPI (blue) to mark nuclei (**A**). Quantifications indicated that disrupting HAUSP by shRNA significantly reduced GSC population (SOX2^+^ or OLIG2+ cells) in GBM xenografts (**B**). Scale bar, 15 µm. Student’s *t* test was used to assess the significance. Data are means ± SD. *n*=3 tumors (200 cells per arm). **, p<0.01. (**C** and **D**) Immunofluorescent staining of the differentiated cell marker GFAP (an astrocyte marker) in GBM xenografts derived from GSCs expressing shHAUSP or shNT (control). Tumor sections were immunostained with a specific antibody against GFAP (in green) and counterstained with DAPI (blue) to mark nuclei (**C**). Quantifications indicated that disrupting HAUSP by shRNA significantly induced cell differentiation in GBM xenografts (**D**). Scale bar, 45 µm. Student’s *t* test was used to assess the significance. Data are means ± SD. *n*=3 tumors (200 cells per arm). **, p<0.001. (**E** and **F**) Immunofluorescent staining of the cell proliferation marker Ki67 in GBM xenografts derived from GSCs expressing shHAUSP or shNT (control). Tumor sections were immunostained with a specific antibody against Ki67 (in red) and counterstained with DAPI (blue) to mark nuclei (**E**). Quantifications indicated that disrupting HAUSP by shRNA significantly reduced cell proliferation in GBM xenografts (**F**). Scale bar, 20 µm. Student’s *t* test was used to assess the significance. Data are means ± SD. *n*=3 tumors (200 cells per arm). ***, p<0.001. (**G** and **H**) TUNEL assay detecting apoptosis in GBM xenografts derived from GSCs expressing shHAUSP or shNT (control). TUNEL assay on tumor sections was performed with an ApopTag Fluorescein *in situ* apoptosis detection kit (in green), and tumor sections were counterstained with DAPI (blue) to mark nuclei (**G**). Quantifications indicated that disrupting HAUSP by shRNA significantly induced cell apoptosis in GBM xenografts (**H**). Scale bar, 45 µm. Student’s *t* test was used to assess significance. Data are means ± SD. *n*=3 tumors (200 cells per arm). ***, p<0.001.

### Pharmacological inhibition of HAUSP suppresses GBM tumor growth and prolongs animal survival

Since GSCs are critical cancer cells to maintain GBM malignant growth and promote therapeutic resistance, disrupting GSCs may effectively improve GBM treatment. As HAUSP stabilizes SOX2 and plays a critical role in maintaining the tumorigenic potential of GSCs, and disrupting HAUSP by shRNA potently inhibits GSC-driven tumor growth, pharmacological targeting of HAUSP could be a promising therapeutic approach to combat GBMs. To test this possibility, we examined whether pharmacological inhibition of HAUSP by its inhibitor P22077 could effectively disrupt GSCs to suppress GBM tumor growth in GSC-derived xenografts. In vitro study showed that P22077 treatment impaired GSC tumorsphere formation in a dose-dependent manner (Figures 7A, S6A and S6B). Detection of the cleaved PARP, an apoptosis marker, by immunoblot analyses confirmed that P22077 treatment potently induced GSC apoptosis in a dose-dependent manner (Figure 7B). Importantly, *in vivo* bioluminescent imaging (IVIS) of intracranial GBM xenografts derived from luciferase-expressing GSCs demonstrated that P22077 treatment (20mg/kg/daily) potently suppressed tumor growth (Figures 7C, 7D, S6C and S6D). As a consequence, P22077 treatment significantly extended the survival of animals bearing the GSC-derived tumors (Figures 7E and S6E). Immunofluorescent analyses of the treated tumors indicated that P22077 treatment markedly reduced SOX2+ cell (GSC) population (Figures 7F, 7G, S6F and S6G), increased cell apoptosis as measured by staining of the activated Caspase 3 (Figures 7H, 7I, S6H and S6I), and inhibited cell proliferation as indicated by Ki67 staining (Figures 7J, 7K, S6J and S6K) in GBM xenografts. We have confirmed the therapeutic impact of targeting HAUSP with P2207 on inhibiting tumor growth in orthotopic xenografts derived from four GSC lines. The potent effects of P22077 on reducing GSCs and inhibiting tumor growth in the intracranial GBM xenografts indicated that P22077 was able to penetrate the blood-brain barrier (BBB) or blood-brain tumor barrier (BTB). Collectively, these data demonstrate that pharmacological inhibition of HAUSP potently suppresses GBM tumor growth, indicating that therapeutic targeting of HAUSP to disrupt GSCs may effectively improve GBM treatment.

**Figure 7.**
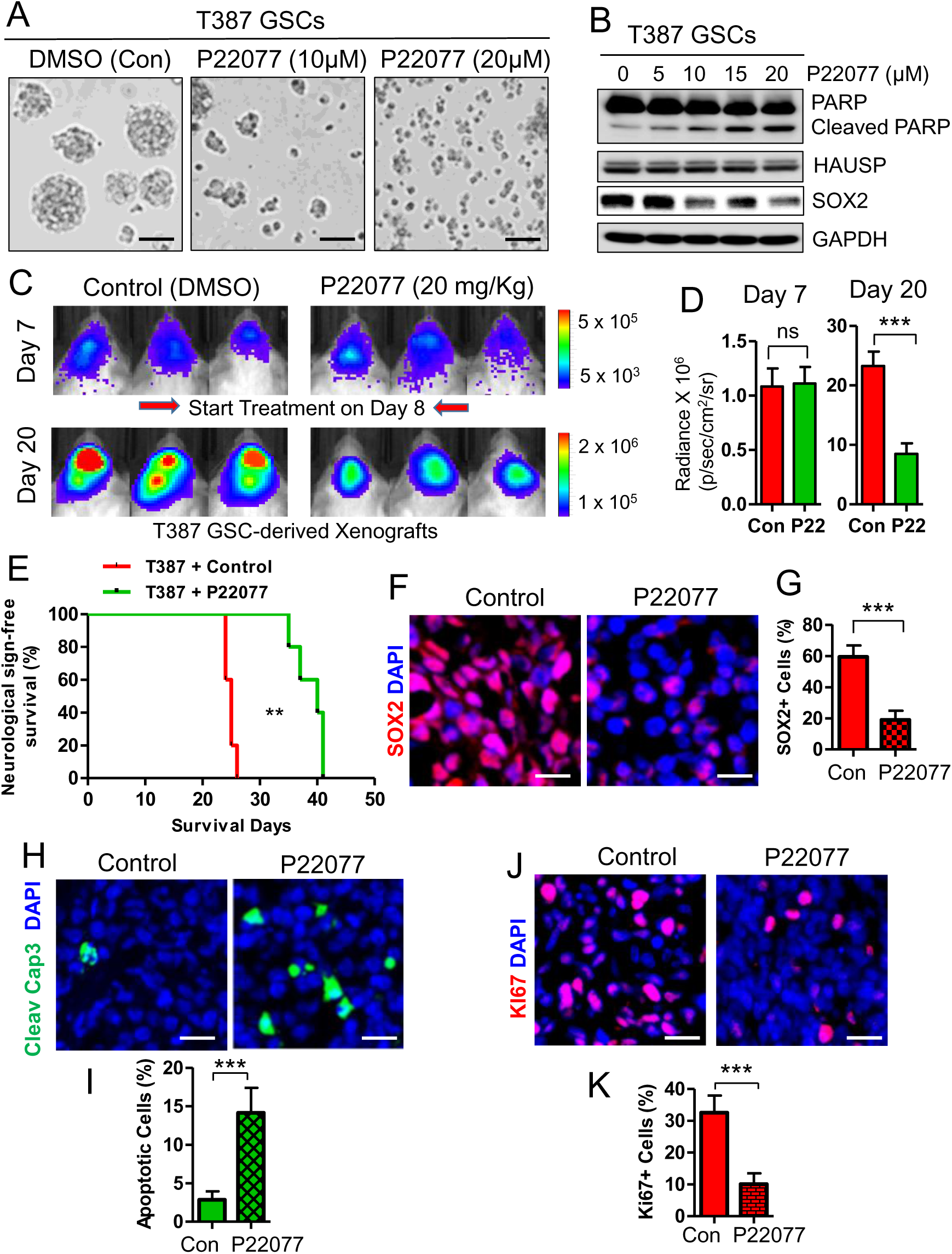
Pharmacological inhibition of HAUSP suppressed GBM tumor growth in GSC-derived xenografts. **(A)** Representative images of tumorspheres derived from GSCs treated with the HAUSP inhibitor P22077 or vehicle control (DMSO). T387 GSCs were treated with P22077 (10 µM or 20 µM) or DMSO for 7 days. Pharmacological inhibition of HAUSP by P22077 reduced GSC tumorsphere formation in a dose-dependent manner. Scale bar, 180 µm. **(B)** Immunoblot analyses of cleaved PARP, total PARP, HAUSP, SOX2 and GAPDH (loading control) in GSCs treated with the increased doses of the HAUSP inhibitor P22077. T387 GSCs were treated with indicated doses (0, 5, 10, 15, 20 µM) of P22077 for 48 hours and then harvested for the immunoblot analyses. (**C** and **D**) *In vivo* bioluminescent imaging (IVIS) of GSC-derived orthotopic xenografts in immunocompromised mice treated with the HAUSP inhibitor P22077 or vehicle control. 7 days after GSC transplantation, the mice were treated with P22077 (20 mg/kg/daily) or vehicle control (DMSO) through tail vein injection. Representative IVIS images (**C**) on day 7 (before treatment) and day 20 (after treatment) are shown. Quantifications of luciferase intensities (**D**) shows that treatment with P22077 significantly inhibited GBM tumor growth in mouse brains. Data are means ± SD. Student’s *t* test was used to assess the significance. *n*=5 mice/group. Ns, no significant difference; ***, P < 0.001. (**E**) Kaplan-Meier survival curves of mice intracranially transplanted with GSCs (T387) and treated with P22077 or vehicle control (DMSO). *n*=5 mice/group. Log-rank analysis was used. **, p<0.01. (**F** and **G**) Immunofluorescent staining of the GSC marker SOX2 in GSC-derived xenografts treated with the HAUSP inhibitor P22077 or vehicle control. Tumor sections were immunostained with a specific antibody against SOX2 (in red) and counterstained with DAPI (blue) to mark nuclei (**F**). Quantification of SOX2+ cells (**G**) indicated that inhibiting HAUSP by P22077 significantly reduced GSC population. Scale bar, 20 µm. Student’s *t* test was used to assess the significance. Data are means ± SD. *n*=3 tumors (200 cells per arm). ***, p<0.001. (**H** and **I**) Immunofluorescent staining of the cell apoptotic marker cleaved caspase 3 in GSC-derived xenografts treated with the HAUSP inhibitor P22077 or vehicle control. Tumor sections were immunostained with a specific antibody against cleaved caspase 3 (in green) and counterstained with DAPI (blue) to mark nuclei (**H**). Quantification of cleaved caspase 3 intensity (**I**) indicated that inhibition of HAUSP with P22077 significantly promoted cell apoptosis in GBM tumors. Scale bar, 20 µm. Student’s *t* test was used to assess significance. Data are means ± SD. *n*=3 tumors (200 cells per arm). ***, p<0.001. (**J** and **K**) Immunofluorescent staining of the cell proliferation marker Ki67 in GSC-derived xenografts treated with P22077 or vehicle control (DMSO). Tumor sections were immunostained with a specific antibody against Ki67 (in red) and counterstained with DAPI (blue) to mark nuclei (**J**). Quantifications of Ki67 signal (**K**) indicated that targeting HAUSP by P22077 significantly reduced cell proliferation in GBM xenografts. Scale bar, 20 µm. Student’s *t* test was used to assess significance. Data are means ± SD. *n*=3 tumors (200 cells per arm). **, p<0.001.

### Targeting HAUSP with P22077 synergizes with radiation to improve therapeutic efficacy for GBM

GBMs are highly resistant to conventional treatments including radiotherapy. Our previous study demonstrated that GSCs are much more resistant to radiation than NSTCs and largely contribute to GBM radioresistance ^16^. Thus, disrupting GSCs may overcome the therapeutic resistance and sensitize GBMs to radiotherapy. As inhibiting HAUSP with its inhibitor P22077 impaired GSC maintenance, promoted GSC differentiation, and potently inhibited GBM tumor growth, we next examined whether targeting HAUSP with P22077 could overcome GSC-mediated radioresistance and sensitize GBM to radiotherapy. *In vivo* bioluminescent imaging analyses demonstrated that the combined treatment with P22077 and irradiation (2 x 2 Gy) indeed showed more effective inhibition on GBM tumor growth relative to the single treatment with P22077 or irradiation (IR) alone in GSC-derived xenografts (Figures 8A-8C and S7A-S7C). As a consequence, the group of mice treated with P22077 in combination with IR survived significantly longer than that treated with P22077 or IR alone (Figures 8D and S7D). Immunofluorescent analyses of cleaved caspase 3 detecting apoptosis on tumor sections showed a significant increase of cell apoptosis in tumors treated with P22077 in combination with IR relative to the tumors with the single treatment (Figures 8E, 8F, S7E and S7F). In addition, the combined IR and P22077 treatment markedly reduced GSC population (SOX2+) relative to IR treatment alone (Figures 8G, 8H, S7G and S7H). Furthermore, Ki67 staining detecting cell proliferation indicated that the P22077 treatment synergized with IR to potently inhibit cell proliferation in GSC-derived xenografts (Figures 8I, 8J, S7I and S7J). Collectively, these data demonstrate that pharmacological inhibition of HAUSP synergizes with radiation and potently suppresses GBM tumor growth to extend animal survival. Our preclinical data indicate that disrupting GSCs through pharmacological inhibition of HAUSP may effectively improve the therapeutic efficacy of radiotherapy for GBM patients.

**Figure 8.**
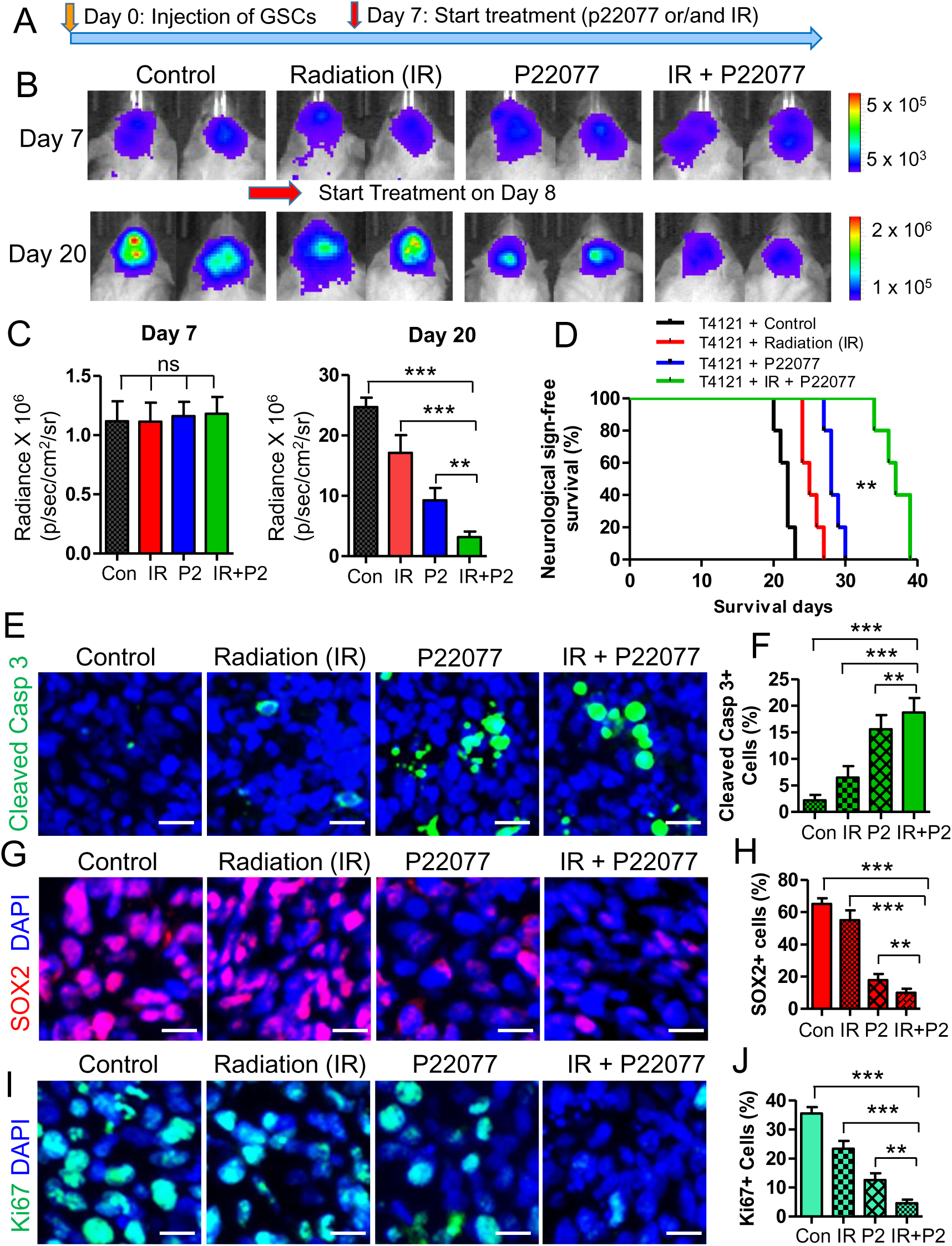
Targeting HAUSP in combination with radiation improved therapeutic efficacy in GSC-derived xenografts. **(A)** A schematic illustration showing the treatment schedule. GSCs (T4121) expressing luciferase were transplanted into brains of immunocompromised NSG mice. 7 days after GSC transplantation, mice were treated with the HAUSP inhibitor P22077 (20 mg/kg/daily) or DMSO (control) through tail vein injection in combination with or without irradiation (IR). IR (2 x 2 Gy) was performed on mouse heads on Day 9 after GSC transplantation in two groups of mice. **(B)** Representative *In vivo* bioluminescent imaging (IVIS) images of intracranial GBM xenografts in the four groups of mice (Control, IR, P22077, IR plus P222077) on Day 7 (before treatment) and Day 20 (after treatment) after GSC transplantation are shown. **(C)** Quantifications of luciferase intensities of intracranial GBM xenografts in the four groups of mice (control, IR, P22077, IR plus P222077) on Day 7 (before treatment) and Day 20 (after treatment) after GSC transplantation. Data are means ± SD. *n*=5. Student’s *t* test was used to assess the significance. **, P < 0.01; ***, p<0.001. **(D)** Kaplan-Meier survival curves of mice intracranially transplanted with GSCs (T4121) and treated with the HAUSP inhibitor P22077, irradiation (IR), P22077 plus IR, or DMSO (control). The combined treatment with P2077 and IR significantly extended the survival of mice bearing the GSC-derived tumors relative the single treatment with P22077 or IR alone. *n*=5 mice per group. P22077 vs control, p<0.003; IR vs control, p<0.01; P22077 plus IR vs control, p<0.001; P22077 vs P22077 plus IR, p<0.01; IR vs P22077 plus IR, p<0.01. Log-rank analysis was used. (**E** and **F**) Immunofluorescent staining of the cell apoptotic marker cleaved caspase 3 in GSC-derived xenografts treated with the HAUSP inhibitor P22077, irradiation (IR), P22077 plus IR, or DMSO (control). Tumor sections were immunostained with a specific antibody against cleaved caspase 3 (in green) and counterstained with DAPI (blue) to mark nuclei (**E**). Quantifications of cleaved caspase 3 intensities (**I**) in GSC-derived xenografts treated with P22077, IR, P22077 plus IR, or DMSO. Scale bar, 15 µm. Student’s *t* test was used to assess the significance. Data are means ± SD. *n*=3 tumors (200 cells per arm). **, p<0.01; ***, p<0.001. (**G** and **H**) Immunofluorescent staining of SOX2 in GSC-derived xenografts treated with the HAUSP inhibitor P22077, irradiation (IR), P22077 plus IR, or DMSO (control). Tumor sections were immunostained with a specific antibody against SOX2 (in red) and counterstained with DAPI (in blue) to mark nuclei (**G**). Quantifications of SOX2+ cells (**H**) in GSC-derived xenografts treated with P22077, IR, P22077 plus IR, or DMSO. Scale bar, 15 µm. Student’s *t* test was used to assess the significance. Data are means ± SD. *n*=3 tumors (200 cells per arm). **, p<0.01; ***, p<0.001. (**I** and **J**) Immunofluorescent staining of the cell proliferation marker Ki67 in GSC-derived xenografts treated with the HAUSP inhibitor P22077, irradiation (IR), P22077 plus IR, or DMSO (control). Tumor sections were immunostained with a specific antibody against Ki67 (in green) and counterstained with DAPI (in blue) to mark nuclei (**I**). Quantifications of Ki67+ cells (**J**) in GSC-derived xenografts treated with P22077, IR, P22077 plus IR, or DMSO. Scale bar, 15 µm. Student’s *t* test was used to assess the significance. Data are means ± SD. *n*=3 tumors (200 cells per arm). **, p<0.01; ***, p<0.001.

## Discussion

Stem cell transcription factors including SOX2 play critical roles in the maintenance of embryonic stem cells (ESc), iPSCs, adult stem cells and cancer stem cells (CSCs). As one of core stem cell transcription factors, SOX2 is preferentially expressed by GSCs in GBMs and play a vital role in the maintenance of GSCs. However, it is very difficult to directly target SOX2. In this study, we demonstrate that HAUSP-mediated deubiquitination and stabilization of SOX2 is required for maintaining the self-renewal and tumorigenic potential of GSCs. Disruption of HAUSP impaired GSC maintenance, induced GSC differentiation, inhibited GSC growth, and suppressed malignant growth, indicating that HAUSP plays an essential role in maintaining GSC properties. Our findings uncover an unappreciated role of HAUSP in the maintenance of GSCs through mediating SOX2 deubiquitination and stabilization to maintain self-renewal and tumorigenic potential of GSCs. Moreover, our preclinical study demonstrated that pharmacological inhibition of HAUSP potently suppressed GBM tumor growth and synergized with radiation to overcome GBM radioresistance, suggesting that targeting HAUSP may effectively improve the therapeutic efficacy to prolong the survival of GBM patients. Thus, our studies identified a promising functional target for developing effective therapeutics targeting GSCs to improve GBM treatment.

Deregulation of the ubiquitination and deubiquitination system has been implicated in the pathogenesis of many diseases, including cancer. USP proteins constituting the largest subfamily of deubiquitinating enzymes (DUBs) usually remove ubiquitin from specific protein substrates and protect these substrates from proteasomal degradation, resulting in stabilization of the proteins or alteration of their localization. HAUSP is one of the most critical DUBs associated with tumorigenesis due to its important role in the negative regulation of p53 and p16^INK4a^ tumor suppressors, as HAUSP deubiquitinates and stabilizes MDM2 (HDM2), the ubiquitin E3 ligase that mediates p53 ubiquitination and degradation ^40, 42^. In this study, we discovered a p53-independent role of HAUSP in the maintenance of GSCs through SOX2 deubiquitination, providing new insights into the molecular mechanisms underlying the HAUSP-mediated tumorigenesis through promoting the self-renewal potential of cancer stem cells. We demonstrate that HAUSP maintains GSC self-renewal and tumorigenic potential through stabilization of SOX2. Targeting HAUSP resulted in SOX2 ubiquitination and proteasomal degradation, thus promoting GSC differentiation and inhibiting cell proliferation. These data indicate that inhibiting HAUSP suppresses GBM tumor growth at least partially through the disruption of GSC maintenance. Notably, targeting HAUSP also differentially upregulates p53 and p21 in GSCs, suggesting that targeting HAUSP may cause double hits to GSCs to inhibit malignant growth. Functional inhibition of HAUSP not only destabilizes SOX2 to induce GSC differentiation and disrupt GSC maintenance but also upregulates p53 and p21 to inhibit cell proliferation. Thus, targeting HAUSP may achieve dual effects to block tumor growth and malignant progression through p53-independent and p53-dependent manners. As HAUSP is a cysteine isopeptidase of the ubiquitin-specific proteases, it is an attractive druggable therapeutic target. In addition, as HAUSP specifically marks GSC population in human primary GBMs, HAUSP may serve as a specific molecular marker of GSCs as well as a useful diagnostic indicator to predict GBM radioresistance.

GBMs are extremely resistant to current treatments including radiation, chemotherapy and immunotherapy ^3–6^. Rapid tumor recurrence after the treatment is a major challenge in the control of GBM malignant progression. Our previous study demonstrated that GSCs promote radioresistance of GBMs by preferential activation of DNA damage checkpoint and repair pathways ^16^. Other studies showed that GSCs are more resistant to chemotherapeutic agents including temozolomide (TMZ) relative to non-stem cancer cells ^19, 24^. GSCs not only directly contribute to the resistance to conventional therapies but also modulate the tumor microenvironment by recruiting and maintaining tumor-supportive macrophages to create an immune suppressive niches ^23, 25, 63^, which promotes malignant progression, therapeutic resistance and thus negatively impacts therapeutic efficacy. In addition, we demonstrated that GSCs not only secrete high levels of VEGF but also generate vascular pericytes to support tumor angiogenesis ^17, 26^. Importantly, GSC-derived tumor pericytes promote the formation of the blood-brain tumor barrier (BTB) that negatively affects chemotherapy by blocking drug delivery ^27^. Thus, disrupting GSCs and CSC-derived pericytes through targeting HAUSP not only overcomes the therapeutic resistance but may also synergizes with other therapies. Although it has been shown that expression levels of HAUSP progressively increase from grade I to IV glioma tumors ^64^, the critical role of HAUSP in maintaining GSC self-renewal and tumorigenic potential through stabilization of SOX2 in GBM has not been reported. Our findings support that HAUSP plays an oncogenic role in GBM malignant growth and progression.

We have demonstrated that pharmacological inhibition of HAUSP by P22077 synergized with radiation to suppress GBM malignant growth and significantly extended the survival of animal bearing GBM tumors. Other study showed that inhibition of USP7/HAUSP with a small molecule inhibitor (P5091) was able to overcome bortezomib resistance and induce apoptosis of multiple myeloma cells ^65^, suggesting that targeting HAUSP may effective solve the resistance problem. Interestingly, a study in neuroblastoma (NB) demonstrated that HAUSP also deubiquitinated and stabilized N-Myc and inhibiting HAUSP effectively inhibited NB tumor growth ^46^. A functional association between HAUSP and DNMT1 has been shown to accelerate oncogenesis and metastasis in colorectal cancer ^41^. In addition, HAUSP deubiquitinates and stabilizes the histone demethylase PHF8, which upregulates a group of genes including cyclin A2 to promote cell growth and proliferation in breast cancer ^47^. As the function of HAUSP has been shown to be associated with malignant progression in several types of cancer including prostate cancer ^44^, breast carcinoma ^47^, leukemia ^49^, epithelial ovarian cancer ^50, 51^, non-small cell lung cancer ^48^, colorectal cancer ^41^, and hepatocellular carcinoma ^45^, targeting HAUSP may inhibit malignant growth of brain metastases from other cancers such as lung cancer. As we demonstrated that targeting HAUSP by P22077 potently suppressed tumor growth of orthotopic GBM in mouse brains, P22077 should be able penetrate the BBB or BTB to inhibit malignant growth of other brain tumors including brain metastases. However, because the solubility of P22077 is not so great, optimizing P22077 or developing more potent HAUSP inhibitors with the capacity to cross the BBB or BTB is urgently needed, which may have great therapeutic potential to improve treatment of highly lethal brain cancers including GBM. Both P22077 and P5091 are covalent catalytic site inhibitors that selectively and covalently modify the catalytic C223 residue to inhibit HAUSP deubiquitinase activity ^58, 59, 66^. Although several allosteric inhibitors including XL188 and its analog XL188 ^67, 68^, FT827 and FT671 ^69, 70^, as well as GNE-6640 and GNE-6676 ^71, 72^ have been developed to target HAUSP, it is not clear whether these inhibitors are able to penetrate the BBB or BTB. It will be highly interesting to examine whether these allosteric inhibitors of HAUSP can also destabilize SOX2 to impair GSC self-renewal and tumorigenic potential.

## Materials and Methods

### Human Glioblastoma (GBM) Specimens and Glioma Stem Cells (GSCs)

De-identified human GBM surgical specimens were collected and provided by the Rose Ella Burkhardt Brain Tumor and Neuro-Oncology Center at Cleveland Clinic in accordance with an Institutional Review Board (IRB)-approved protocol. Informed consents were obtained from all patients under the surgery. Glioma stem cells (GSCs) and matched non-stem tumor cells (NSTCs) were isolated from primary GBMs or patient-derived GBM xenografts (PDXs) and cultured as previously described ^16, 26, 27, 61, 62^. Briefly, total glioma cells were isolated from GBM tumors with the Papain Dissociation System (Worthington Biochemical) according to the protocol provided by the manufacturer. After recovery of the isolated glioma cells in the stem cell medium (Neurobasal-A medium with B27 supplement, 20 ng/ml EGF and 20 ng/ml bFGF) for 6-8 hours to allow the re-expression of GSC surface markers, then the cells were labeled with a FITC-conjugated anti-CD15 antibody (Millipore, CBL144F) and a phycoerythrin (PE)-conjugated anti-CD133 antibody (Miltenyi Biotec, 130-098-826) followed by the fluorescence-activated cell sorting (FACS) to sort the GSC population (CD15+/CD133+) and the matched NSTC population (CD15-/CD133-). The sorted GSCs were validated by three functional assays, including self-renewal (in vitro serial neurosphere formation), multipotent differentiation (induction of multi-lineage differentiation *in vitro*), and tumor initiation and formation (*in vivo* limiting dilution assay) to confirm the cancer stem cell property as previously described ^16, 26, 27, 62^. The validated GSCs were then used for the in vitro and in vivo experiments.

### Establishment of Intracranial GBM Xenografts, in vivo Treatments and Bioluminescent Imaging

All animal procedures were performed in accordance with a protocol approved by the Cleveland Clinic Institutional Animal Care and Use Committee (IACUC). Establishment of orthotopic GBM xenografts through intracranial transplantation of GSCs was performed as previously described ^16, 27, 61, 62^. GSCs were transduced with luciferase and/or HAUSP shRNA (shHAUSP) or non-targeting shRNA control (shNT) through lentiviral infection, and then selected with puromycin (1 µg/mL, Fisher Scientific) for 48 hours after the infection. GSCs were transplanted into immunocompromised NSG mice (Jackson Laboratories or the Cleveland Clinic’s Biological Resources Unit) through intracranial injection. Intracranial tumor growth in animals was monitored by in vivo bioluminescent imaging (IVIS) before and after treatment, and maintained until manifestation of neurological signs. For IVIS, mice were treated with D-luciferin (1.5 mg/10g, LUCK-GOLDBIO) through intraperitoneal injection and anesthetized with isoflurane before imaging. Luciferase images of intracranial tumors were captured under an IVIS imaging system (Spectrum CT, PerkinElmer). To examine the effect of HAUSP inhibition with P22077 (BOC Sciences, B0084-462597) on tumor growth, mice were treated with P22077 (20mg/Kg/daily) or vehicle control through tail vein injection. For the irradiation (IR), mice bearing GSC-derived intracranial GBM tumors were irradiated (2 Gy) for 2-5 times in the Pantak X-ray irradiator at the Cleveland Clinic Lerner Research Institute. Mouse brains bearing GBM tumors were collected for further analyses including immunofluorescent and immunohistochemical staining.

### Immunoblot Analysis and Immunoprecipitation

Immunoblot analyses were performed as previously described ^60–62^. Briefly, GSCs or NSTCs were lysed in the cell lysis buffer (1% TritonX-100, 10% glycerol, 50mM HEPES pH7.5, 150mM NaCl, 100mM NaF, 1mM PMSF, 1mM Na3VO4, protease inhibitor cocktail). Cellular proteins were separated by SDS-PAGE and transferred to PVDF membrane (Biorad), then blocked by 5% milk for 30 minutes, and incubated with a primary antibody overnight at 4°C. Membranes were washed with TBST for 3 times, incubated with a secondary antibody for 2 hours at room temperature, washed with TBST for 3 times, and then subjected to chemiluminescent substrate (Thermo Scientific). Signals were detected with ChemiDoc XRS+ Imager (Biorad).

For immunoprecipitation (IP), cells lysates were incubated with 20 μL of Protein A/G agarose gel (Santa Cruz, sc-2003) and a primary antibody with constant rotation overnight at 4 °C. The IP complexes were washed with ice-cold 0.3% Triton X-100 in PBS buffer for three times and eluted in loading buffer by boiling for 10 min, and then analyzed by immunoblot analyses. Proteins were resolved on NuPAGE Novex 4-12% Bis-Tris gels (Invitrogen, NP0322BOX), transferred to PVDF membranes, and immunoblotted with indicated antibodies.

Specific antibodies against HAUSP (Bethyl, A300-033A), SOX2 (Bethyl, A201-741A; or Santa Cruz, SC-365964), OLIG2 (R&D System, AF2418), GFAP (Biolegend, 644702), TUJ1 (Biolegend, 801201), MAP2 (Biolegend, 801801), GAPDH (R&D System, MAB5718), HA (Sigma, 11867423001), α-Tubulin (Sigma, T6199), Flag (Sigma, F1804), Ubiquitin (Biolegend, 646302), Cleaved Caspase 3 (Cell Signaling, 9661), Cleaved PARP (Cell Signaling, 5625), and anti-Flag-agarose (Sigma, A2220) were used for immunoblot and/or immunoprecipitation.

### Mass Spectrometry

To identify potential deubiquitinase(s) for SOX2 in GSCs, SOX2-interacting proteins were pulled down from GSCs by immunoprecipitation and then analyzed by Mass Spectrometry. GSCs were transduced with Flag-tagged-SOX2 or a vector control, and the cell lysates were co-immunoprecipitated by anti-Flag agarose (Sigma, A2220), separated by SDS-PAGE and visualized by staining. The SDS-PAGE gel was dehydrated in acetonitrile, and the precipitated proteins were digested with trypsin. The peptides were then analyzed and identified through Liquid Chromatography Mass Spectrometry (LC-MS) by the Proteomics Core at Cleveland Clinic Lerner Research Institute.

### Ubiquitination assay

The ubiquitination assay in vitro or in vivo was performed as previously described ^53, 60^. Cells were treated with MG132 (20 µM) for 6-8 hours before harvest, then lysed in the IP buffer (50 mM Hepes, 150 mM NaCl, 1% NP-40, and 5 mM EDTA) in the presence of phosphatase and protease inhibitors (4906837001 and 4693159001, Roche). Cell lysate was subjected to immunoprecipitation with the related antibody and agarose beads, and the product was eluted in loading buffer by boiling for 10 min. The precipitated proteins were resolved on NuPAGE Novex 4-12% Bis-Tris gels (Invitrogen, NP0322BOX), blotted onto PVDF membranes, and probed by the anti-ubiquitin antibody.

### Immunofluorescent and Immunohistochemical Staining

Immunofluorescent staining were performed as previously described ^26, 27, 62, 73^. Human GBM surgical specimens, intracranial GBM xenografts or GSC tumorspheres were fixed with 4% PFA overnight at 4°C, stored in 30% sucrose solution overnight at 4°C, embedded in OCT overnight at -20°C, and cryosectioned at a thickness of 8 microns. Tumor sections were blocked with 1% BSA (Sigma) in a PBS solution containing 0.03% TWEEN-20 for 1 hour at RT and incubated with a primary antibody (1:200 dilution) overnight at 4°C. Specific antibodies against HAUSP (Bethyl, A300-033A), SOX2 (Bethyl, A201-741A; or Santa Cruz, sc-17320), Olig2 (R&D systems AF2418), the EC marker CD31 (DAKO, M082301), GFAP (Biolegend, 840001), TUJ1 (Biolegend, 801201), the cell apoptotic marker cleaved caspase-3 (Cell Signaling, 9661), and the cell proliferation marker Ki67 (Cell Signaling, 9129) were used for the immunofluorescent staining on GBM tumor sections, tumorsphere sections, or fixed cells cultured on coverslips. TUNEL Assay detecting apoptotic cell death were performed with the ApopTag Fluorescein *in situ* apoptosis detection kit (S7110, Millipore) according to the manufacturer’s instructions. Immunohistochemical (IHC) staining on sections of primary tumors and normal tissues were performed with an ABC kit and the 3, 3’-diaminobezine detection kit (Vector Laboratories) as previously described ^23, 62^. A specific anti-HAUSP antibody (Bethyl, IHC-00018) was used for the IHC staining on GBM tumor sections.

### Differentiation Assay

To assess the differentiation potential of sorted GSCs or examine the effect of HAUSP disruption on GSC differentiation, isolated GSCs or GSCs expressing shHAUSP were cultured on plates or coverslips pretreated with the Matrigel (354230; BD Biosciences) and induced for differentiation in serum-containing medium (1% FBS in DME) or in Neurobasal medium without EGF and FGF growth factors but with B27 (12587-010, Invitrogen). Differentiated cells or control GSCs were collected for immunoblot analyses or fixed with PFA for immunofluorescent staining at the indicated days as described above.

### Plasmid Constructs and Lentivirus Production

Lentivirus constructs expressing shHAUSP (TRCN0000004057 or TRCN0000004059) or shNT (SHC002) were obtained from Sigma-Aldrich. The lentiviral plasmid expressing wild type HAUSP (WT) or catalytically dead HAUSP (HAUSP-CS) was constructed by inserting human HAUSP-WT or HAUSP-CS ORF cDNA into the pCDH-CMV-MCS-EF1a-Puro vector (System Biosciences, CD510B-1). The lentiviral plasmid expressing SOX2 was constructed by cloning human SOX2 ORF cDNA into the pCDH-CMV-MCS-EF1a-Puro vector (System Biosciences, CD510B-1) or PCW57.1 vector (Addgene, 50661). For lentivirus production, 293FT cells were transduced with the desired plasmid together with the helper plasmids pCI-VSVG and ps-PAX2. 2-3 days after the transfection, lentiviral supernatant was collected and titered as described previously ^27, 62, 73^. GSCs were then infected with lentiviruses at a MOI around 3.

### Quantitative RT-PCR

Total mRNA was isolated from GSCs with the RNeasy kit (QIAGEN) and reversely transcribed into cDNA with the PrimeScript RT Master Mix (Takara Bio., Inc.). Quantitative RT-PCR was performed with the SYBR green master mix (Applied Biosystems, 4309155) in the real time PCR system (Applied Biosystems). Data were from at least three independent experiments. The following primers were used to quantify mRNA of genes: HAUSP forward primer: 5’-ACT TTG AGC CAC AGC CCG GTA ATA-3’, HAUSP reverse primer: 5’-GCC TTG AAC ACA CCA GCT TGG AAA-3’; OLIG2 forward primer: 5’-CAA GAA GCA AAT GAC AGA GCC GGA-3’, OLIG2 reverse primer: 5’-TGG TGA GCA TGA GGA TGT AGT TGC-3’; GAPDH forward primer: 5’-TGT TGC CAT CAA TGA CCC CTT CA-3’, GAPDH reverse primer: 5’-CTC CAC GAC GTA CTC AGC GCC-3’.

### Quantification and Statistical Analysis

All data were analyzed by using the GraphPad Prism 8 (GraphPad Inc.). The two-tailed unpaired Student’s t test was used to assess the difference between two groups, and the one-way analysis of variance (ANOVA) was used to compare differences among more than two groups, with p<0.05 being considered statistically significant. Analyses of animal survival were performed using the Kaplan-Meier curve method, with the log-rank test for comparison. All quantitative data are means ± SD or means ± SEM as indicated in the figure legend.

## Acknowledgements

We thank the Rose Ella Burkhardt Brain Tumor and Neuro-Oncology Center of Cleveland Clinic for providing GBM surgical specimens for this study. We appreciate the assistance provided by the Proteomics, Imaging, Cell Culture, Genomic and Flow Cytometry Cores of the Lerner Research Institute. This research work was supported by the Cleveland Clinic Foundation and NIH R01 grants (CA184090, NS091080 and NS099175) to S. Bao, and NIH Shared Instrument Grants (S10OD018205 and 1S10RR031537) to the Cleveland Clinic Lerner Research Institute.

## Author Contributions

S.B. and Z.H. developed the scientific concept and working hypothesis, and designed the experimental approaches. Z.H. and S. B. analyzed the data and prepared the manuscript. Z.H., K.Z., W.T., X.F., Q.H. and Q.W. performed the experiments. J.L., J.S.Y. and J.N.R. provided scientific input for the manuscript.

The authors declare no competing financial interests.

## References

1. Furnari, F.B. et al. Malignant astrocytic glioma: genetics, biology, and paths to treatment. Genes Dev 21, 2683–2710 (2007).

2. Stupp, R. et al. Radiotherapy plus concomitant and adjuvant temozolomide for glioblastoma. N Engl J Med 352, 987–996 (2005).

3. Zhang, H. et al. Glioblastoma Treatment Modalities besides Surgery. J Cancer 10, 4793–4806 (2019).

4. Wen, P.Y. & Kesari, S. Malignant gliomas in adults. N Engl J Med 359, 492–507 (2008).

5. Stupp, R. et al. Effects of radiotherapy with concomitant and adjuvant temozolomide versus radiotherapy alone on survival in glioblastoma in a randomised phase III study: 5-year analysis of the EORTC-NCIC trial. Lancet Oncol 10, 459–466 (2009).

6. Reardon, D.A. et al. Effect of Nivolumab vs Bevacizumab in Patients With Recurrent Glioblastoma: The CheckMate 143 Phase 3 Randomized Clinical Trial. JAMA Oncol 6, 1003–1010 (2020).

7. Magee, J.A., Piskounova, E. & Morrison, S.J. Cancer stem cells: impact, heterogeneity, and uncertainty. Cancer Cell 21, 283–296 (2012).

8. Medema, J.P. Cancer stem cells: the challenges ahead. Nat Cell Biol 15, 338–344 (2013).

9. Valent, P. et al. Cancer stem cell definitions and terminology: the devil is in the details. Nat Rev Cancer 12, 767–775 (2012).

10. Galli, R. et al. Isolation and characterization of tumorigenic, stem-like neural precursors from human glioblastoma. Cancer Res 64, 7011–7021 (2004).

11. Hemmati, H.D. et al. Cancerous stem cells can arise from pediatric brain tumors. Proc Natl Acad Sci U S A 100, 15178–15183 (2003).

12. Singh, S.K. et al. Identification of human brain tumour initiating cells. Nature 432, 396–401 (2004).

13. Prager, B.C., Bhargava, S., Mahadev, V., Hubert, C.G. & Rich, J.N. Glioblastoma Stem Cells: Driving Resilience through Chaos. Trends Cancer 6, 223–235 (2020).

14. Huang, Z., Cheng, L., Guryanova, O.A., Wu, Q. & Bao, S. Cancer stem cells in glioblastoma--molecular signaling and therapeutic targeting. Protein Cell 1, 638–655 (2010).

15. Lathia, J.D., Mack, S.C., Mulkearns-Hubert, E.E., Valentim, C.L. & Rich, J.N. Cancer stem cells in glioblastoma. Genes Dev 29, 1203–1217 (2015).

16. Bao, S. et al. Glioma stem cells promote radioresistance by preferential activation of the DNA damage response. Nature 444, 756–760 (2006).

17. Bao, S. et al. Stem cell-like glioma cells promote tumor angiogenesis through vascular endothelial growth factor. Cancer Res 66, 7843–7848 (2006).

18. Cheng, L. et al. Elevated invasive potential of glioblastoma stem cells. Biochem Biophys Res Commun 406, 643–648 (2011).

19. Liu, G. et al. Analysis of gene expression and chemoresistance of CD133+ cancer stem cells in glioblastoma. Mol Cancer 5, 67 (2006).

20. Calabrese, C. et al. A perivascular niche for brain tumor stem cells. Cancer Cell 11, 69–82 (2007).

21. Li, Z. et al. Hypoxia-inducible factors regulate tumorigenic capacity of glioma stem cells. Cancer Cell 15, 501–513 (2009).

22. Wei, J. et al. Glioma-associated cancer-initiating cells induce immunosuppression. Clinical cancer research : an official journal of the American Association for Cancer Research 16, 461–473 (2010).

23. Zhou, W. et al. Periostin secreted by glioblastoma stem cells recruits M2 tumour-associated macrophages and promotes malignant growth. Nat Cell Biol 17, 170–182 (2015).

24. Zheng, H. et al. p53 and Pten control neural and glioma stem/progenitor cell renewal and differentiation. Nature 455, 1129–1133 (2008).

25. Tao, W. et al. Dual Role of WISP1 in maintaining glioma stem cells and tumor-supportive macrophages in glioblastoma. Nat Commun 11, 3015 (2020).

26. Cheng, L. et al. Glioblastoma stem cells generate vascular pericytes to support vessel function and tumor growth. Cell 153, 139–152 (2013).

27. Zhou, W. et al. Targeting Glioma Stem Cell-Derived Pericytes Disrupts the Blood-Tumor Barrier and Improves Chemotherapeutic Efficacy. Cell stem cell 21, 591–603 e594 (2017).

28. Masui, S. et al. Pluripotency governed by Sox2 via regulation of Oct3/4 expression in mouse embryonic stem cells. Nat Cell Biol 9, 625–635 (2007).

29. Boumahdi, S. et al. SOX2 controls tumour initiation and cancer stem-cell functions in squamous-cell carcinoma. Nature 511, 246–250 (2014).

30. Kim, J.B. et al. Pluripotent stem cells induced from adult neural stem cells by reprogramming with two factors. Nature 454, 646–650 (2008).

31. Kamachi, Y. & Kondoh, H. Sox proteins: regulators of cell fate specification and differentiation. Development 140, 4129–4144 (2013).

32. Mu, P. et al. SOX2 promotes lineage plasticity and antiandrogen resistance in TP53- and RB1-deficient prostate cancer. Science 355, 84–88 (2017).

33. Basu-Roy, U. et al. Sox2 antagonizes the Hippo pathway to maintain stemness in cancer cells. Nat Commun 6, 6411 (2015).

34. Ferone, G. et al. SOX2 Is the Determining Oncogenic Switch in Promoting Lung Squamous Cell Carcinoma from Different Cells of Origin. Cancer Cell 30, 519–532 (2016).

35. Bass, A.J. et al. SOX2 is an amplified lineage-survival oncogene in lung and esophageal squamous cell carcinomas. Nat Genet 41, 1238–1242 (2009).

36. Suva, M.L. et al. Reconstructing and reprogramming the tumor-propagating potential of glioblastoma stem-like cells. Cell 157, 580–594 (2014).

37. Fang, L. et al. A methylation-phosphorylation switch determines Sox2 stability and function in ESC maintenance or differentiation. Mol Cell 55, 537–551 (2014).

38. Holowaty, M.N. et al. Protein profiling with Epstein-Barr nuclear antigen-1 reveals an interaction with the herpesvirus-associated ubiquitin-specific protease HAUSP/USP7. J Biol Chem 278, 29987–29994 (2003).

39. Boutell, C., Canning, M., Orr, A. & Everett, R.D. Reciprocal activities between herpes simplex virus type 1 regulatory protein ICP0, a ubiquitin E3 ligase, and ubiquitin-specific protease USP7. J Virol 79, 12342–12354 (2005).

40. Cummins, J.M. et al. Tumour suppression: disruption of HAUSP gene stabilizes p53. Nature 428, 1 p following 486 (2004).

41. Du, Z. et al. DNMT1 stability is regulated by proteins coordinating deubiquitination and acetylation-driven ubiquitination. Sci Signal 3, ra80 (2010).

42. Li, M., Brooks, C.L., Kon, N. & Gu, W. A dynamic role of HAUSP in the p53-Mdm2 pathway. Mol Cell 13, 879–886 (2004).

43. Li, M. et al. Deubiquitination of p53 by HAUSP is an important pathway for p53 stabilization. Nature 416, 648–653 (2002).

44. Song, M.S. et al. The deubiquitinylation and localization of PTEN are regulated by a HAUSP-PML network. Nature 455, 813–817 (2008).

45. Cai, J.B. et al. Ubiquitin-specific protease 7 accelerates p14(ARF) degradation by deubiquitinating thyroid hormone receptor-interacting protein 12 and promotes hepatocellular carcinoma progression. Hepatology 61, 1603–1614 (2015).

46. Tavana, O. et al. HAUSP deubiquitinates and stabilizes N-Myc in neuroblastoma. Nat Med 22, 1180–1186 (2016).

47. Wang, Q. et al. Stabilization of histone demethylase PHF8 by USP7 promotes breast carcinogenesis. J Clin Invest 126, 2205–2220 (2016).

48. Zhao, G.Y. et al. USP7 overexpression predicts a poor prognosis in lung squamous cell carcinoma and large cell carcinoma. Tumour Biol 36, 1721–1729 (2015).

49. Carra, G. et al. Therapeutic inhibition of USP7-PTEN network in chronic lymphocytic leukemia: a strategy to overcome TP53 mutated/deleted clones. Oncotarget 8, 35508–35522 (2017).

50. Ma, M. & Yu, N. Ubiquitin-specific protease 7 expression is a prognostic factor in epithelial ovarian cancer and correlates with lymph node metastasis. Onco Targets Ther 9, 1559–1569 (2016).

51. Zhang, L., Wang, H., Tian, L. & Li, H. Expression of USP7 and MARCH7 Is Correlated with Poor Prognosis in Epithelial Ovarian Cancer. Tohoku J Exp Med 239, 165–175 (2016).

52. Sheng, Y. et al. Molecular recognition of p53 and MDM2 by USP7/HAUSP. Nat Struct Mol Biol 13, 285–291 (2006).

53. Huang, Z. et al. Deubiquitylase HAUSP stabilizes REST and promotes maintenance of neural progenitor cells. Nat Cell Biol 13, 142–152 (2011).

54. van der Horst, A. et al. FOXO4 transcriptional activity is regulated by monoubiquitination and USP7/HAUSP. Nat Cell Biol 8, 1064–1073 (2006).

55. Faustrup, H., Bekker-Jensen, S., Bartek, J., Lukas, J. & Mailand, N. USP7 counteracts SCFbetaTrCP- but not APCCdh1-mediated proteolysis of Claspin. J Cell Biol 184, 13–19 (2009).

56. Kon, N. et al. Roles of HAUSP-mediated p53 regulation in central nervous system development. Cell Death Differ 18, 1366–1375 (2011).

57. Kon, N. et al. Inactivation of HAUSP in vivo modulates p53 function. Oncogene 29, 1270–1279 (2010).

58. Altun, M. et al. Activity-based chemical proteomics accelerates inhibitor development for deubiquitylating enzymes. Chem Biol 18, 1401–1412 (2011).

59. Pozhidaeva, A. et al. USP7-Specific Inhibitors Target and Modify the Enzyme’s Active Site via Distinct Chemical Mechanisms. Cell Chem Biol 24, 1501–1512 e1505 (2017).

60. Fang, X. et al. Deubiquitinase USP13 maintains glioblastoma stem cells by antagonizing FBXL14-mediated Myc ubiquitination. J Exp Med 214, 245–267 (2017).

61. Shi, Y. et al. Ibrutinib inactivates BMX-STAT3 in glioma stem cells to impair malignant growth and radioresistance. Sci Transl Med 10 (2018).

62. Guryanova, O.A. et al. Nonreceptor tyrosine kinase BMX maintains self-renewal and tumorigenic potential of glioblastoma stem cells by activating STAT3. Cancer Cell 19, 498–511 (2011).

63. Shi, Y. et al. Tumour-associated macrophages secrete pleiotrophin to promote PTPRZ1 signalling in glioblastoma stem cells for tumour growth. Nat Commun 8, 15080 (2017).

64. Cheng, C., Niu, C., Yang, Y., Wang, Y. & Lu, M. Expression of HAUSP in gliomas correlates with disease progression and survival of patients. Oncol Rep 29, 1730–1736 (2013).

65. Chauhan, D. et al. A small molecule inhibitor of ubiquitin-specific protease-7 induces apoptosis in multiple myeloma cells and overcomes bortezomib resistance. Cancer Cell 22, 345–358 (2012).

66. Weinstock, J. et al. Selective Dual Inhibitors of the Cancer-Related Deubiquitylating Proteases USP7 and USP47. ACS Med Chem Lett 3, 789–792 (2012).

67. Schauer, N.J. et al. Selective USP7 inhibition elicits cancer cell killing through a p53-dependent mechanism. Sci Rep 10, 5324 (2020).

68. Lamberto, I. et al. Structure-Guided Development of a Potent and Selective Non-covalent Active-Site Inhibitor of USP7. Cell Chem Biol 24, 1490–1500 e1411 (2017).

69. Turnbull, A.P. et al. Molecular basis of USP7 inhibition by selective small-molecule inhibitors. Nature 550, 481–486 (2017).

70. Gavory, G. et al. Discovery and characterization of highly potent and selective allosteric USP7 inhibitors. Nat Chem Biol 14, 118–125 (2018).

71. Di Lello, P. et al. Discovery of Small-Molecule Inhibitors of Ubiquitin Specific Protease 7 (USP7) Using Integrated NMR and in Silico Techniques. J Med Chem 60, 10056–10070 (2017).

72. Kategaya, L. et al. USP7 small-molecule inhibitors interfere with ubiquitin binding. Nature 550, 534–538 (2017).

73. Zhang, A. et al. Protein Sumoylation with SUMO1 Promoted by Pin1 in Glioma Stem Cells Augments Glioblastoma Malignancy. Neuro Oncol (2020).

