## Supplemental Figures and Legends for "HAUSP Stabilizes SOX2 through Deubiquitination to Maintain Self-renewal and Tumorigenic Potential of Glioma Stem Cells"

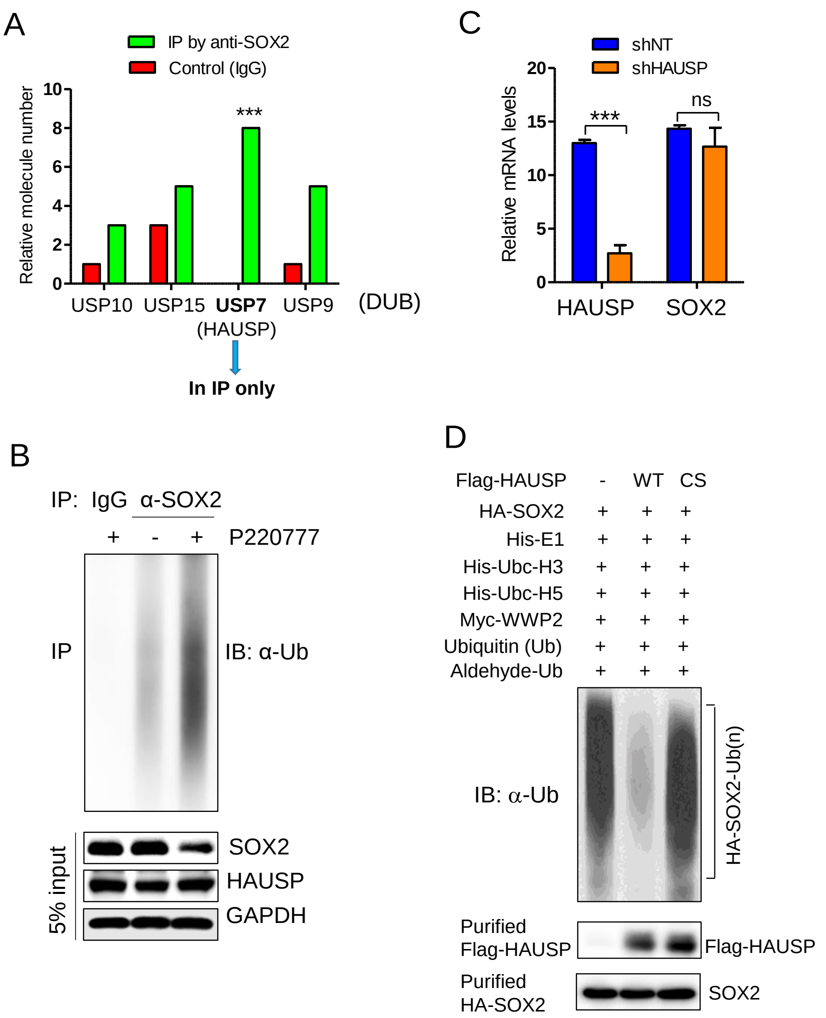


**Supplemental Figure S1. Identification of HAUSP as the key deubiquitinase of SOX2 in GSCs.**

(**A**) Identification of potential deubiquitinase(s) that interacted with SOX2 in GSCs through immunoprecipitation (IP) and Mass Spectrometric analyses. T4121 GSCs were transduced with Flag-SOX2, and the cell lysates were co-immunoprecipitated with anti-Flag agarose or IgG (control). IP complexes were separated by SDS-PAGE, digested with trypsin, and analyzed through Liquid Chromatography Mass Spectrometry (LC-MS). Potential deubiquitinases including USP7 (HAUSP), USP9, USP10 and USP15 were identified in the SOX2 IP complex, but HAUSP is the dominant deubiquitinase only present in the SOX2 IP complex but not in the control (IgG)

IP complex.

(**B**) Ubiquitination assay showing the effect of HAUSP inhibition by P22077 on poly-ubiquitination of SOX2 in GSCs. T4121 GSCs were treated with P22077 (20 μM) or vehicle control (DMSO) for 48 hours, then treated with MG132 for 6 hours, and harvested for immunoprecipitation (IP) with anti-SOX2 antibody or IgG control. The IP products were immunoblotted with anti-ubiquitin (Ub) antibody to assess the SOX2 ubiquitination in GSCs with or without HAUSP inhibition. 5% cell lysates were immunoblotted with antibodies against SOX2, HAUSP and GAPDH (control).

(**C**) Quantitative RT-PCR analyses of relative mRNA levels of HAUSP and SOX2 in GSCs transduced with shHAUSP or shNT control. GSCs (T4121) were transduced with HAUSP shRNA (shHAUSP) or non-targeting shRNA (shNT) through lentiviral infection for 48 hours, and total RNAs were extracted for RT-PCR analyses. Student t test was used to assess the significance. ns, no significant difference; ***, $p<0.001$.

(**D**) In vitro ubiquitination assay to examine the effect of wild-type HAUSP (Flag-HAUSP-WT) or catalytically dead mutant HAUSP (Flag-HAUSP-CS) on poly-ubiquitination of HA-SOX2. Purified Flag-HAUSP-WT or Flag-HAUSP-CS were incubated with purified HA-SOX2 in an in vitro ubiquitination system, and resulting products were immunoblotted with anti-ubiquitin (Ub) antibody to assess the HA-SOX2 ubiquitination in presence of wild-type HAUSP or the mutant HAUSP.

Huang et al. Fig. S2

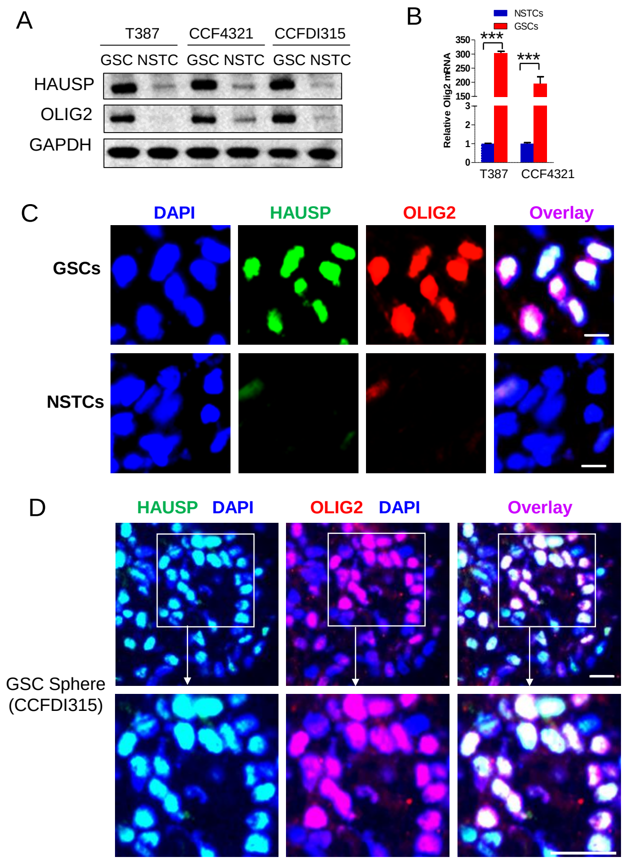


**Supplemental Figure S2. HAUSP is preferentially expressed in glioma stem cells (GSCs) isolated from human primary GBMs.**

(**A**) Immunoblot analyses of HAUSP and OLIG2 (a GSC marker) in GSCs and matched non-stem tumor cells (NSTCs) isolated from four human GBM tumors. HAUSP is preferentially expressed in GSCs expressing OLIG2 relative to NSTCs.

(**B**) Quantitative RT-PCR analyses of HAUSP expression in two pairs of GSCs and matched NSTCs. HAUSP mRNA is differentially expressed in GSCs relative to NSTCs. Student t test was used to assess the significance. ***, $p<0.001$.

(**C**) Immunofluorescent staining of HAUSP and the GSC marker OLIG2 in GSCs and matched NSTCs isolated from primary a GBM (CCF3264). GSCs and matched NSTCs were immunostained with specific antibodies against HAUSP (in green) and OLIG2 (in red), and then counterstained with DAPI to indicate nuclei (in blue). Scale bar, 10 μm.

(**D**) Immunofluorescent staining of HAUSP and OLIG2 in GSC tumorsphere. Sections of GSC tumorspheres (CCFDI315) were immunostained with specific antibodies against HAUSP (in green) and OLIG2 (in red), and then counterstained with DAPI to indicate nuclei (in blue). A small portion of the GSC sphere indicated by the white square was enlarged and shown in the bottom panels. Scale bar, 20 μm.

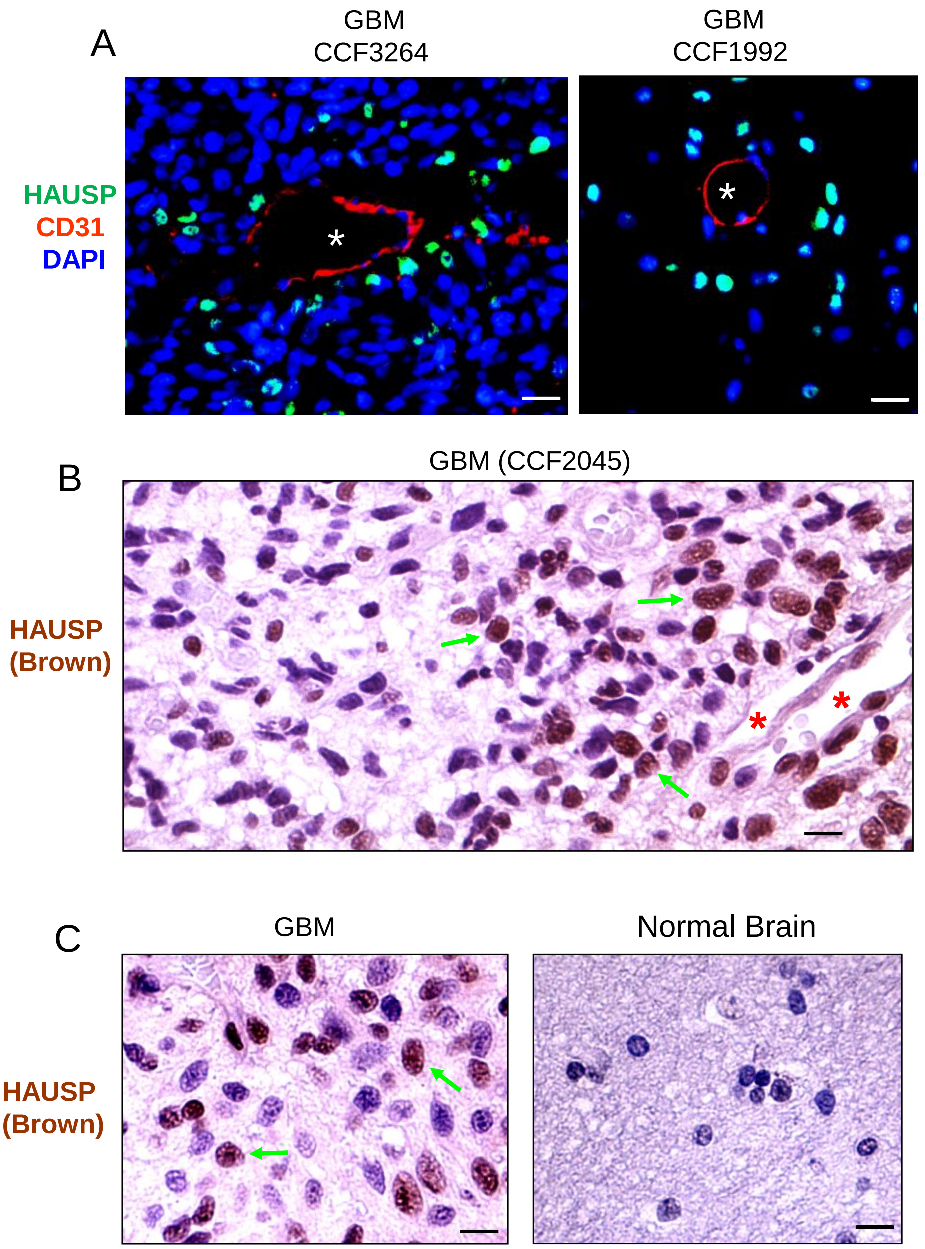


**Supplemental Figure S3. HAUSP and SOX2 are co-expressed in GSCs in perivascular niches in human primary GBMs.**

(**A**) Immunofluorescent staining of HAUSP and the endothelial cell marker CD31 in human primary GBMs. Frozen sections of GBM (CCF3264 or CCF1992) were immunostained with specific antibodies against HAUSP (in green) and CD31 (in red), and then counterstained with DAPI to indicate nuclei (in blue). White stars indicate vessel lumens in GBM tumors. Scale bar, 20 μm.

(**B** and **C**) Immunohistochemical (IHC) staining of HAUSP in human primary GBM and normal brain tissue. Paraffin sections of a primary GBM (CCF2045) and a normal brain tissue were immunostained with a specific antibody against HAUSP (in brown), and counterstained with hematoxylin to indicate nuclei. HAUSP (in brown) is expressed by some glioma cells around the vessel (indicated by stars) in GBM tumors (**B, C**), but HAUSP is rarely expressed in the normal brain tissue (**C**). Scale bar, 15 μm.

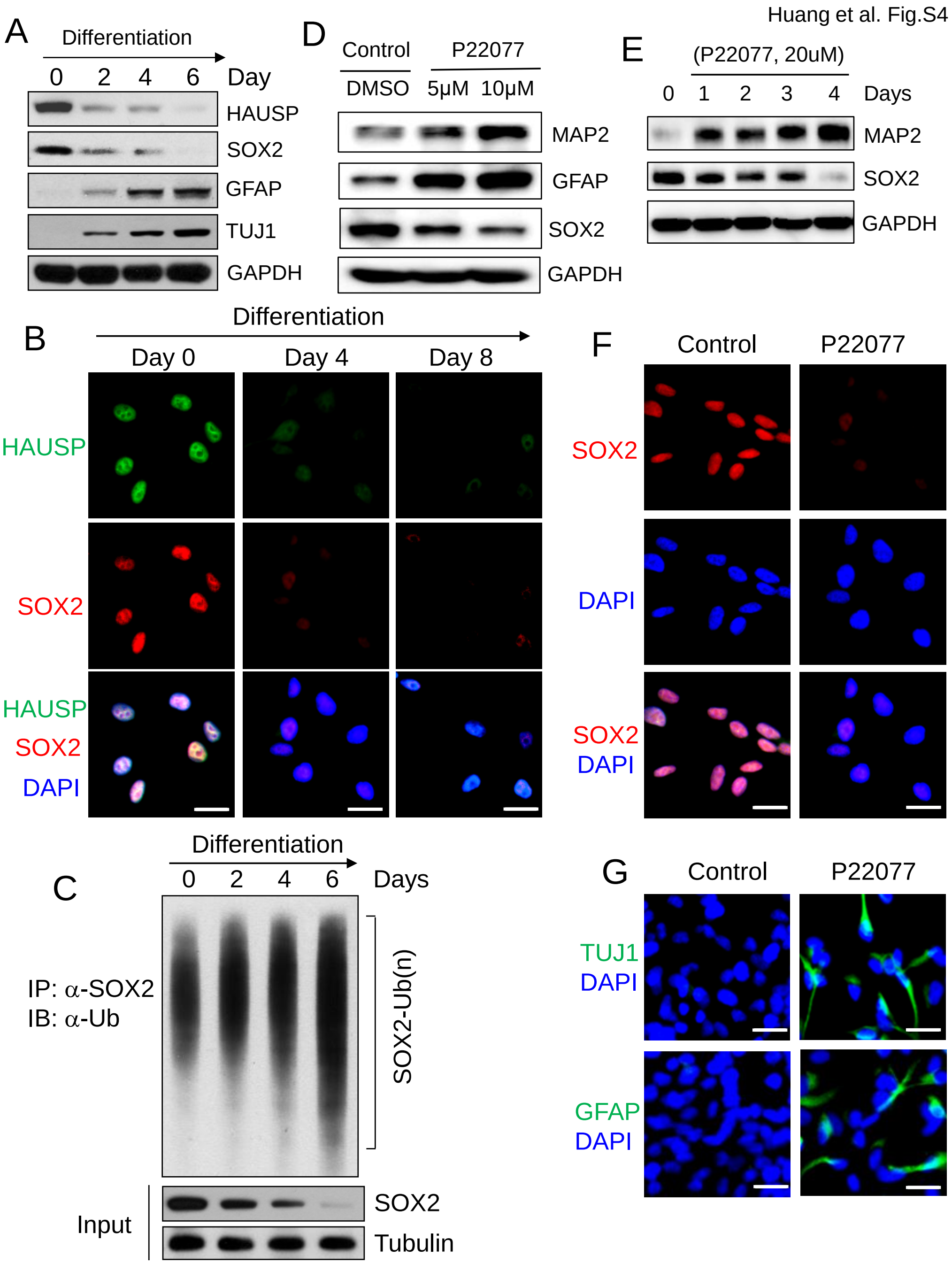


**Supplemental Figure S4. HAUSP levels were reduced during GSC differentiation and inhibiting HAUSP by P22077 promoted GSC differentiation.**

(**A**) Immunoblot analyses of HAUSP, SOX2, GFAP (an astrocyte marker) and TUJ1 (a neuronal marker) during GSC differentiation induced by serum. T387 GSCs were induced by serum (5%) for differentiation and harvested at the indicated days for the analyses.

(**B**) Immunofluorescent staining of HAUSP and SOX2 during GSC differentiation induced by serum. T387 GSCs were induced by serum (5%) for differentiation and harvested at the indicated days for the staining with specific antibodies against HAUSP (in green) and SOX2 (in red). Cells were counterstained with DAPI to mark nuclei (in blue). Scale bar, 25 μm.

(**C**) Ubiquitination assay detecting poly-ubiquitination of SOX2 (SOX2-Ub) during GSC differentiation. T387 GSCs were induced by serum (5%) for differentiation for the indicated days, then treated with MG132 for 6 hours, and harvested for immunoprecipitation (IP) with anti-SOX2 antibody. The IP products were immunoblotted with anti-ubiquitin (Ub) antibody to assess the SOX2 ubiquitination during GSC differentiation. 5% cell lysates were immunoblotted with antibodies against SOX2 and Tubulin (control).

(**D**) Immunoblot analyses of GFAP (an astrocyte marker), MAP2 (a neuronal marker) and SOX2 in GSCs treated with the HAUSP inhibitor P22077 or vehicle control (DMSO). T387 GSCs were treated with P22077 (5 or 10μM) or DMSO for 3 days and then harvested for the analyses.

(**E**) Immunoblot analyses of MAP2 (a neuronal marker) and SOX2 in GSCs treated with the HAUSP inhibitor P22077 for increased times. T387 GSCs were treated with P22077 (20 μM) or vehicle control (DMSO) for 1-4 days and then harvested for the analyses.

(**F**) Immunofluorescent staining of SOX2 in GSCs treated with the HAUSP inhibitor P22077 or vehicle control. T387 GSCs were treated with P22077 (20 μM) or vehicle control (DMSO) for 3 days and then harvested for the staining with a specific antibody against SOX2 (in red). Cells were counterstained with DAPI to mark nuclei (in blue). Scale bar, 25 μm.

(**G**) Immunofluorescent staining of the differentiated cell marker TUJ1 or GFAP in GSCs treated with the HAUSP inhibitor P22077 or vehicle control. T387 GSCs were treated with P22077 (10 μM) or vehicle control (DMSO) for 4 days and then harvested for the staining with a specific antibody against TUJ1 or GFAP (in green). Cells were counterstained with DAPI to mark nuclei (in blue). Scale bar, 25 μm.

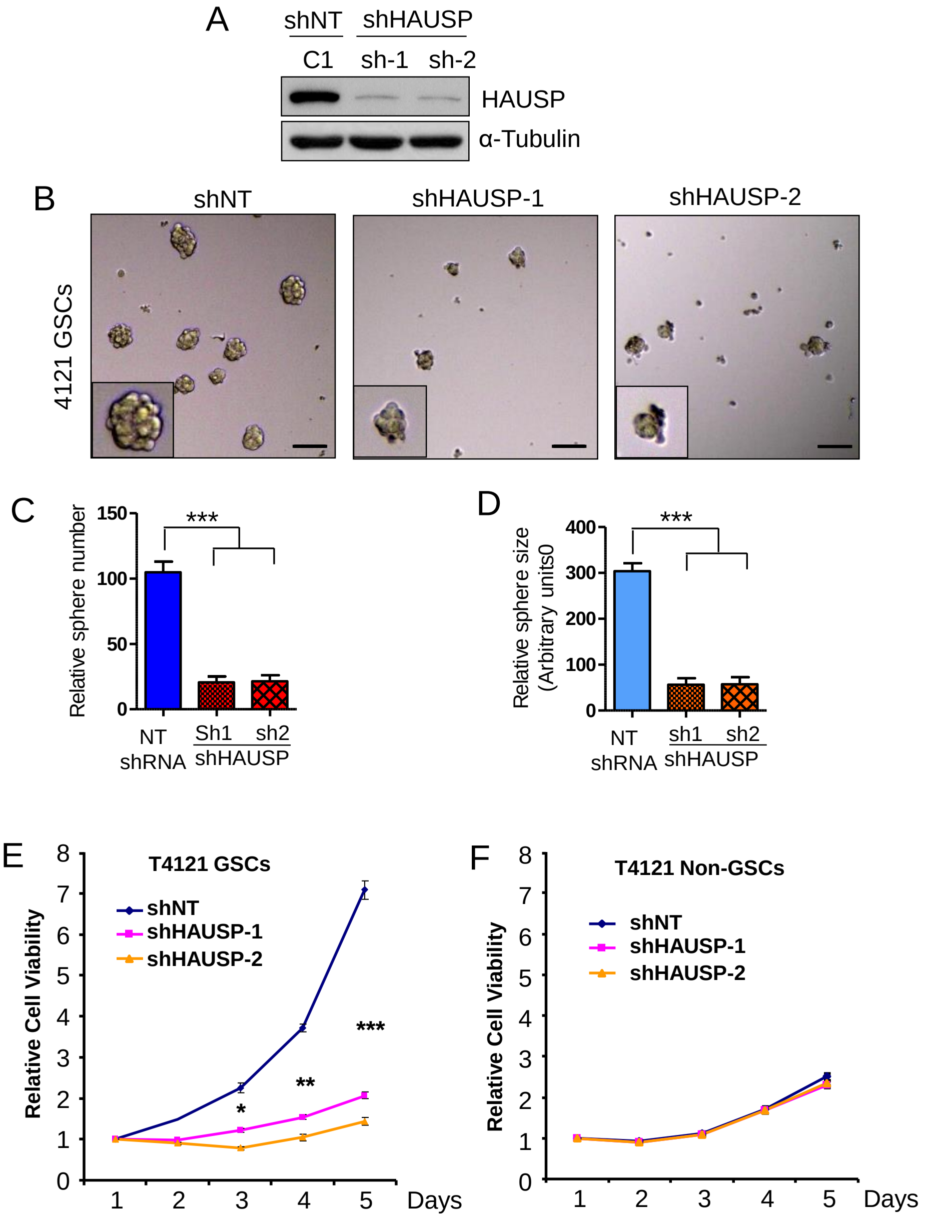


**Supplemental Figure S5. Disrupting HAUSP inhibited GSC tumorsphere formation and cells proliferation.**

(**A**) Immunoblot analysis of HAUSP in GSCs transduced with shHAUSP or shNT (control). T4121 GSCs were infected with shHAUSP (sh-1 or sh-2) or shNT lentiviruses for 48 hours and then harvested for the analysis.

(**B-D**) Tumorsphere formation of GSCs transduced with shHAUSP or shNT control. T4121 GSCs were transduced with shHAUSP (shHAUSP-1 or shHAUSP-2) or shNT through lentiviral infection for 48 hours and cultured for 6 days in the Neurobasal medium with B27 and EGF/bFGF. Representative images of GSC tumorspheres are shown (**B**). Quantifications of relative tumorsphere number (**C**) and size (**D**) showing that HAUSP knockdown significantly reduced GSC tumorsphere formation. Scale bar, 250 μm. ANOVA analysis was used to assess the significance. ***, p<0.001.

(**E** and **F**) Cell viability assay of GSCs (**E**) and matched NSTCs (**F**) transduced with shHAUSP or shNT (control). T4121 GSCs and matched NSTCs were infected with shHAUSP (shHAUSP-1 or shHAUSP-2) or shNT lentiviruses, and cell viability of GSCs or NSTCs were measured from day 1 through day 5. Student t test was used to assess the significance. *, p<0.05; **, p<0.01, ***, p<0.001.

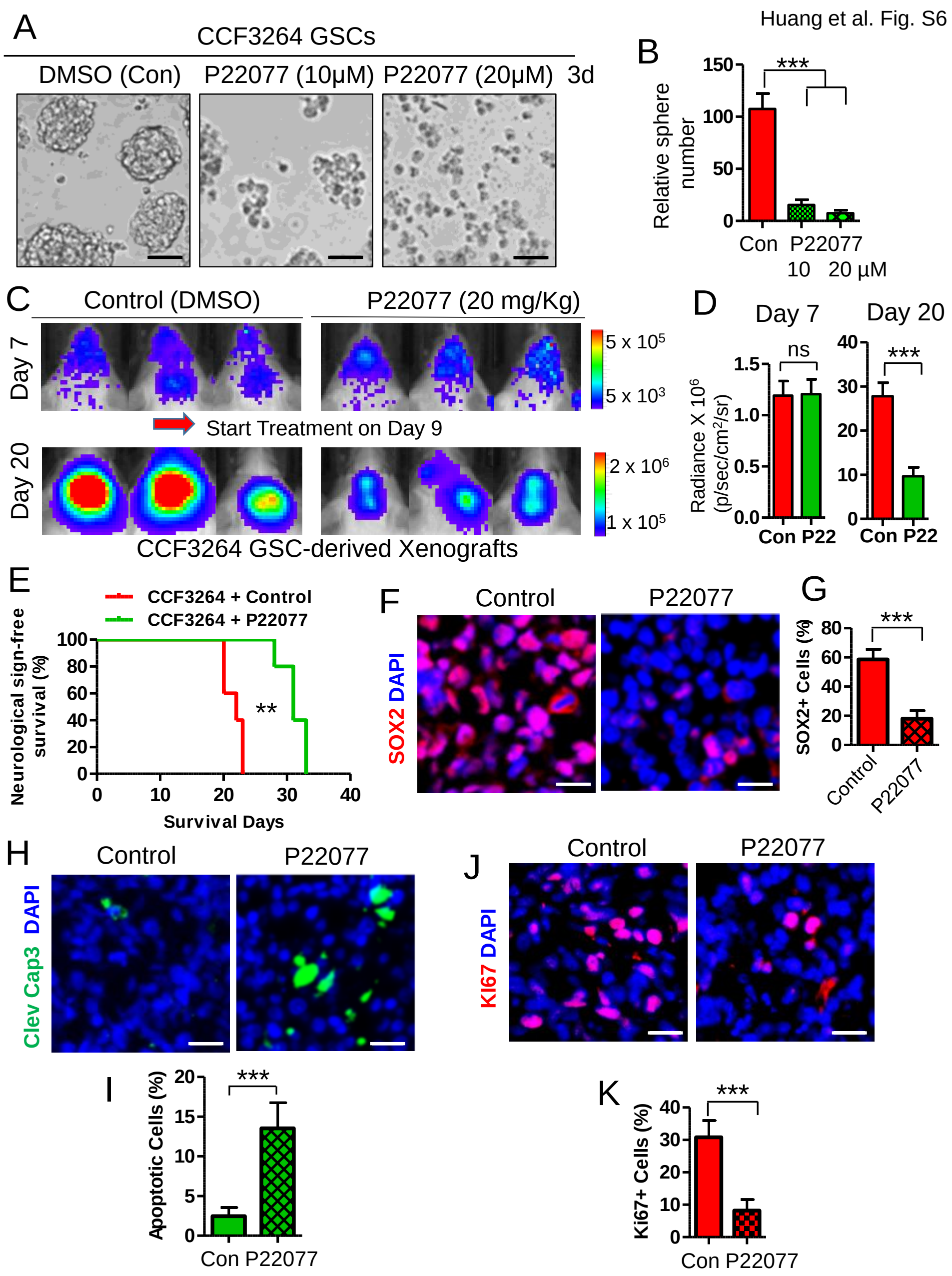


**Supplemental Figure S6. Pharmacological inhibition of HAUSP suppressed GBM tumor growth in GSC-derived xenografts.**

(**A**) Representative images of tumorspheres derived from GSCs treated with the HAUSP inhibitor P22077 or vehicle control (DMSO). GSCs (CCF3264) were treated with P22077 (10 μM or 20 μM) or DMSO for 7 days. Pharmacological inhibition of HAUSP by P22077 reduced GSC tumorsphere formation in a dose-dependent manner. Scale bar, 160 μm.

(**B**) Quantifications of relative numbers of tumorspheres derived from GSCs treated with P22077. GSCs (CCF3264) were treated with P22077 (10 or 20 μM) or vehicle control (DMSO) for 7 days, and the relative numbers of GSC tumorspheres were analyzed. ANOVA analysis was used to assess the significance. ***, p<0.001.

(**C** and **D**) *In vivo* bioluminescent imaging (IVIS) of GSC-derived orthotopic xenografts in immunocompromised mice treated with the HAUSP inhibitor P22077. GSCs (CCF3264) were transduced with luciferase and transplanted into NSG mice through intracranial injection. 7 days after GSC transplantation, the mice were treated with P22077 (20 mg/kg/daily) or vehicle control (DMSO) through tail vein injection. Representative IVIS images at day 7 (before treatment) and day 20 (after treatment) are shown (**C**). Quantifications of luciferase intensities shows that treatment with P22077 significantly inhibited GBM tumor growth in mouse brains (**D**). Data are means ± SD. Student's *t* test was used to assess the significance. *n*=5 mice/group. Ns, no significant difference; ***, P < 0.001.

(**E**) Kaplan-Meier survival curves of mice intracranially transplanted with GSCs (CCF3264) and treated with P22077 or vehicle control (DMSO). *n*=5 mice/group. Log-rank analysis was used. **, p<0.01.

(**F** and **G**) Immunofluorescent staining of the GSC marker SOX2 in GSC-derived xenografts treated with the HAUSP inhibitor P22077 or vehicle control. Tumor sections were immunostained with a specific antibody against SOX2 (in red) and counterstained with DAPI (blue) to mark nuclei (**F**). Quantifications indicated that inhibiting HAUSP by P22077 significantly reduced $SOX2^+$ cell population (GSCs) (**G**). Scale bar, 20 μm. Student's *t* test was used to assess the significance. Data are means ± SD. *n*=3 tumors (200 cells per arm). ***, p<0.001.

(**H** and **I**) Immunofluorescent staining of the cell apoptotic marker cleaved caspase 3 in GSC-derived xenografts treated with the HAUSP inhibitor P22077 or vehicle control. Tumor sections were immunostained with a specific antibody against cleaved caspase 3 (in green) and counterstained with DAPI (blue) to mark nuclei (**H**). Quantification indicated that inhibition of HAUSP with P22077 significantly promoted cell apoptosis in GBM tumors (**I**). Scale bar, 20 μm. Student's *t* test was used to assess the significance. Data are means ± SD. *n*=3 tumors (200 cells per arm). ***, p<0.001.

(**J** and **K**) Immunofluorescent staining of the cell proliferation marker Ki67 in GSC-derived xenografts treated with P22077 or vehicle control (DMSO). Tumor sections were immunostained with a specific antibody against Ki67 (in red) and counterstained with DAPI (blue) to mark nuclei (J). Quantifications indicated that targeting HAUSP by P22077 significantly reduced cell proliferation in GBM xenografts (K). Scale bar, 20 μm. Student's *t* test was used to assess the significance. Data are means ± SD. *n*=3 tumors (200 cells per arm). ***, p<0.001.

Huang et al. Fig. S7

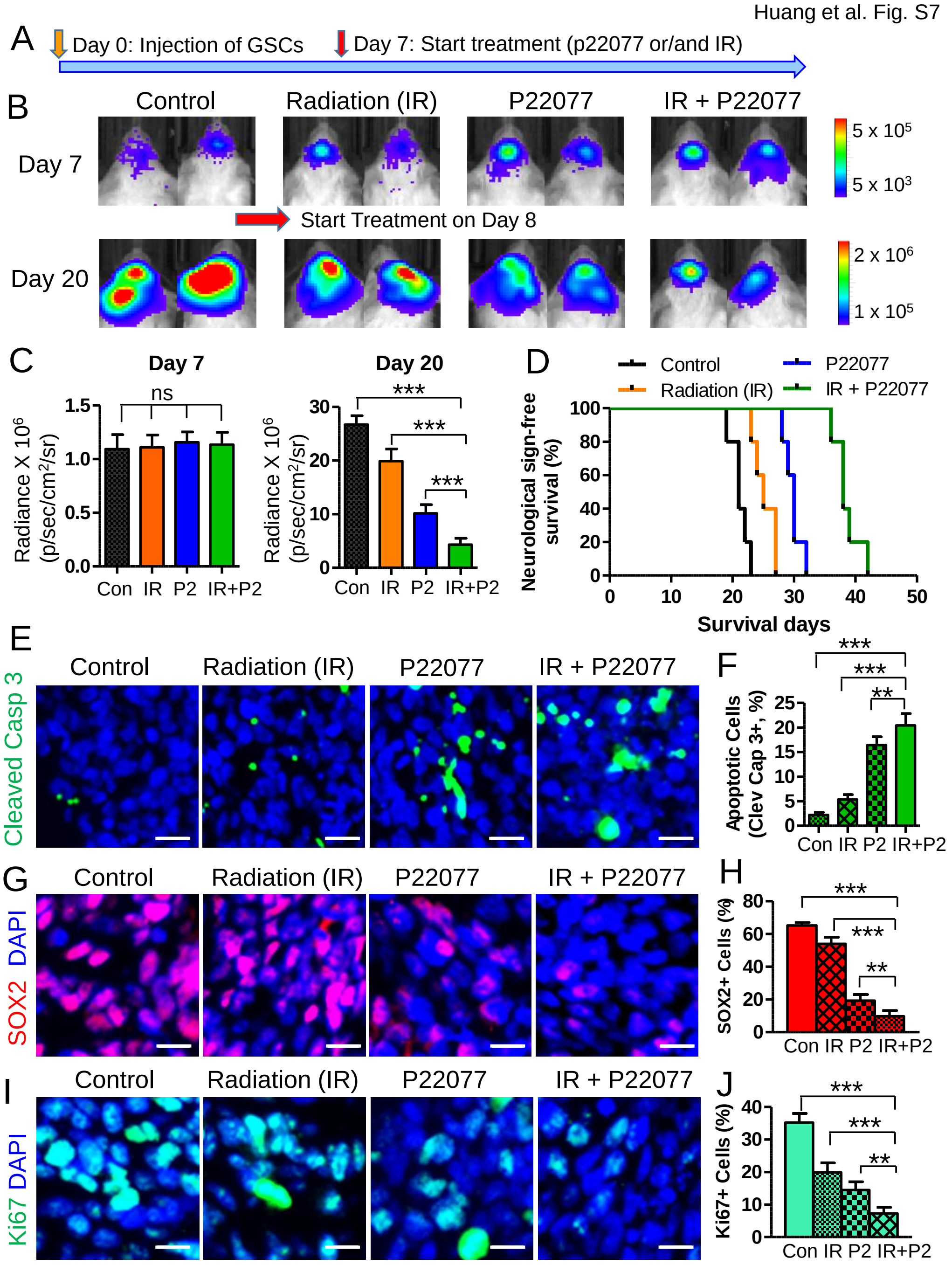


**Supplemental Figure S7. Targeting HAUSP in combination with radiation improved therapeutic efficacy in GSC-derived xenografts.**

(**A**) A schematic illustration showing the treatment schedule. GSCs (CCF3264) expressing luciferase were transplanted into brains of immunocompromised NSG mice. 7 days after GSC transplantation, mice were treated with the HAUSP inhibitor P22077 (20 mg/kg/daily) or DMSO (control) through tail vein injection in combination with or without irradiation (IR). IR (2 x 2 Gy) was performed on mouse heads on Day 9 after GSC transplantation in two groups of mice.
